# High Throughput Virtual Screening and Validation of a SARS-CoV-2 Main Protease Non-Covalent Inhibitor

**DOI:** 10.1101/2021.03.27.437323

**Authors:** Austin Clyde, Stephanie Galanie, Daniel W. Kneller, Heng Ma, Yadu Babuji, Ben Blaiszik, Alexander Brace, Thomas Brettin, Kyle Chard, Ryan Chard, Leighton Coates, Ian Foster, Darin Hauner, Vilmos Kertesz, Neeraj Kumar, Hyungro Lee, Zhuozhao Li, Andre Merzky, Jurgen G. Schmidt, Li Tan, Mikhail Titov, Anda Trifan, Matteo Turilli, Hubertus Van Dam, Srinivas C. Chennubhotla, Shantenu Jha, Andrey Kovalevsky, Arvind Ramanathan, Martha S. Head, Rick Stevens

**Affiliations:** Data Science and Learning Division, Argonne National Laboratory, Lemont, IL 60439, USA; Department of Computer Science, University of Chicago, Chicago, IL 60615, USA; Neutron Scattering Division, Oak Ridge National Laboratory, Oak Ridge, TN, 37831, USA; Biosciences Division, Oak Ridge National Laboratory, Oak Ridge, TN, 37831 USA; Department of Electrical and Computer Engineering, Rutgers University, Piscataway, NJ 08854, USA; Computational Sciences Initiative, Brookhaven National Laboratory, Upton, New York 11973, USA; Bioscience Division, Los Alamos National Laboratory, Los Alamos, New Mexico 87545, USA; Consortium for Advanced Science and Engineering, University of Chicago, Chicago, IL 60615, USA; Computing Environment and Life Sciences Directorate, Argonne National Laboratory, Lemont, IL 60439, USA; Second Target Station, Oak Ridge National Laboratory, Oak Ridge, TN 37831, USA; Globus, University of Chicago, Chicago, IL 60615, USA; National Virtual Biotechnology Laboratory, US Department of Energy, Washington DC, USA; University of Illinois at Urbana-Champaign, Champaign, IL 61820, USA; Joint Institute for Biological Sciences, Oak Ridge National Laboratory, Oak Ridge, TN, 37831 USA; Department of Computational and Systems Biology, University of Pittsburgh, Pittsburgh, PA, 15260 USA; Computational Biology Group, Biological Science Division, Pacific Northwest National Laboratory, Richland, WA 99352, United States

**Keywords:** Room-temperature X-ray crystallography, Molecular dynamics, Machine learning, Virtual Screening

## Abstract

Despite the recent availability of vaccines against the acute respiratory syndrome coronavirus 2 (SARS-CoV-2), the search for inhibitory therapeutic agents has assumed importance especially in the context of emerging new viral variants. In this paper, we describe the discovery of a novel non-covalent small-molecule inhibitor, MCULE-5948770040, that binds to and inhibits the SARS-Cov-2 main protease (M^pro^) by employing a scalable high throughput virtual screening (HTVS) framework and a targeted compound library of over 6.5 million molecules that could be readily ordered and purchased. Our HTVS framework leverages the U.S. supercomputing infrastructure achieving nearly 91% resource utilization and nearly 126 million docking calculations per hour. Downstream biochemical assays validate this M^pro^ inhibitor with an inhibition constant (*K*_*i*_) of 2.9 *µ*M [95% CI 2.2, 4.0]. Further, using room-temperature X-ray crystallography, we show that MCULE-5948770040 binds to a cleft in the primary binding site of M^pro^ forming stable hydrogen bond and hydrophobic interactions. We then used multiple *µ*s-timescale molecular dynamics (MD) simulations, and machine learning (ML) techniques to elucidate how the bound ligand alters the conformational states accessed by M^pro^, involving motions both proximal and distal to the binding site. Together, our results demonstrate how MCULE-5948770040 inhibits M^pro^ and offers a springboard for further therapeutic design.

Significance Statement
The ongoing novel coronavirus pandemic (COVID-19) has prompted a global race towards finding effective therapeutics that can target the various viral proteins. Despite many virtual screening campaigns in development, the discovery of validated inhibitors for SARS-CoV-2 protein targets has been limited. We discover a novel inhibitor against the SARS-CoV-2 main protease. Our integrated platform applies downstream biochemical assays, X-ray crystallography, and atomistic simulations to obtain a comprehensive characterization of its inhibitory mechanism. Inhibiting M^pro^ can lead to significant biomedical advances in targeting SARS-CoV-2 treatment, as it plays a crucial role in viral replication.

## 1. Introduction

The ongoing novel coronavirus pandemic (COVID-19) has resulted in over 100 million infections and more than 2 million deaths worldwide^*^. Although vaccines against the COVID-19 causative agent, the severe acute respiratory syndrome coronavirus 2 (SARS-CoV-2), are being deployed (1, 2), the discovery of drugs which can inhibit various SARS-CoV-2 proteins remains essential for treating patients (3, 4). Leveraging existing coronavirus treatments developed for severe acute respiratory syndrome (SARS) and middle eastern respiratory syndrome (MERS) (5), as well as broad international collaborations, researchers have quickly determined structures for over 15 viral proteins, including inhibitor/lead bound structures and fragment-based screening for several non-structural proteins (NSP) such as the main protease (3C-like protease/M^pro^), adenine diphosphate ribosyl-transferase (ADRP/NSP3), endoribonuclease (NSP15), and helicase (NSP13) (6), all playing crucial roles in viral replication. Together, these collaborations have significantly accelerated the design and development of antiviral treatments targeting SARS-CoV-2 (7, 8).

Of these proteins M^pro^ is an attractive drug target mainly because it plays a critical role in viral replication and does not have any closely related homologs within the human genome (9, 10). Drug discovery efforts have resulted in discovering/re-purposing small molecules based on their ability to inhibit other coronavirus M^pro^ from middle east respiratory syndrome(MERS) and severe acute respiratory syndrome(SARS); however, it has been a challenge to identify non-covalent inhibitors for SARS-CoV-2 M^pro^ mainly due to the intrinsic flexibility of the primary binding site (11).

High throughput virtual screening (HTVS) is a common step of drug-discovery, enabling rapid, low-cost screening of significantly larger compound libraries than feasible in experimental studies (12). A number of efforts have focused on creating open HTVS infrastructure, taking advantage of cloud computing platforms or supercomputing resources to support large-scale ligand docking across various protein targets (13). These platforms have leveraged open-source toolkits such as AutoDock/AutoDock-VINA (for molecular docking) (14) in conjunction with molecular modeling (MM) and molecular dynamics (MD) simulation engines to capture ‘modes’ of interaction between a protein target and specific compounds from compound-libraries (e.g., ZINC (15), MCULE (16)). Of these approaches, the COVID-Moonshot project, using crowdsourced design strategies, high-throughput experimental screening, MD simulations and ML were able to identify both covalent and non-covalent inhibitors against M^pro^ which demonstrated viral inhibition *in vitro* (17).

In this paper, we describe our discovery of a non-covalent small molecule inhibitor for M^pro^ using our HTVS platform that employs super-computing resources, ensemble docking strategies, high-throughput experimental screening, X-ray crystallography, and MD. Complementary to efforts that scaled crowdsourcing approaches (18) as well as HTVS across potentially *O*(billion) compounds (13, 19), we used a library consisting of 6.5 million in-stock compounds from the MCULE library (16). Ensemble docking was carried out across available crystal structures for M^pro^, from which *O*(1,000) top consensus-scoring compounds across two popular docking programs (Autodock-VINA (14) and OpenEye FRED (20)) were experimentally characterized. From these compounds, we discovered one molecule that inhibits M^pro^ with K_*i*_ of 2.9 *µ*M and determined its room-temperature X-ray crystallographic structure to 1.8 Å resolution. Finally, we used *µ*s-timescale atomistic MD simulations to characterize the binding mechanism to the M^pro^ active site, while altering the enzyme’s overall conformational dynamics. Our workflow provides a scalable framework for the rapid discovery of viable lead molecules against SARS-CoV-2. All of our generated data, including ensemble docking results and MD simulations, are freely available and can be used for future studies.

## 2. Results

### A. HTVS of M^pro^ with on-demand molecular libraries

A docking screen against the main protease of SARS-CoV2, M^pro^, was performed on an orderable on-demand compound library from Mcule (16). Given the intrinsic flexibility of the M^pro^’s primary binding pocket consisting of the four conserved binding sites (S1’, S1, S2 and S4) (11, 21), we used five different crystal structures were used for an ensemble docking approach using PDB identifiers 6LU7(22), 6W63 (23), 7BQY (22), 7C7P (24), and 7JU7 (25). In addition to the structural ensemble, we used the docking protocols and scoring functions from the OpenEye Scientific FRED (20) toolkit. In total, over 63 million docking scores were computed over the five structures, two compound libraries, and two protocols. The overall workflow is summarized in Fig. 1a. The workflows were deployed on HPC resources at the Argonne and Oak Ridge leadership computing facilities (ALCF/OLCF) using Theta and Summit supercomputers and using the Texas Advanced Computing Center (TACC; Frontera) and San Diego Supercomputing Center (SDSC; Comet). The resulting docking libraries (including scripts of preparation and docking) and the docking scores are available as a downloadable dataset (26). The details of the computational performance and workflow optimization are described in the Supporting Information (SI) text (sections S1 and S2) and Fig. S1(a)-(c).

**Fig. 1.**
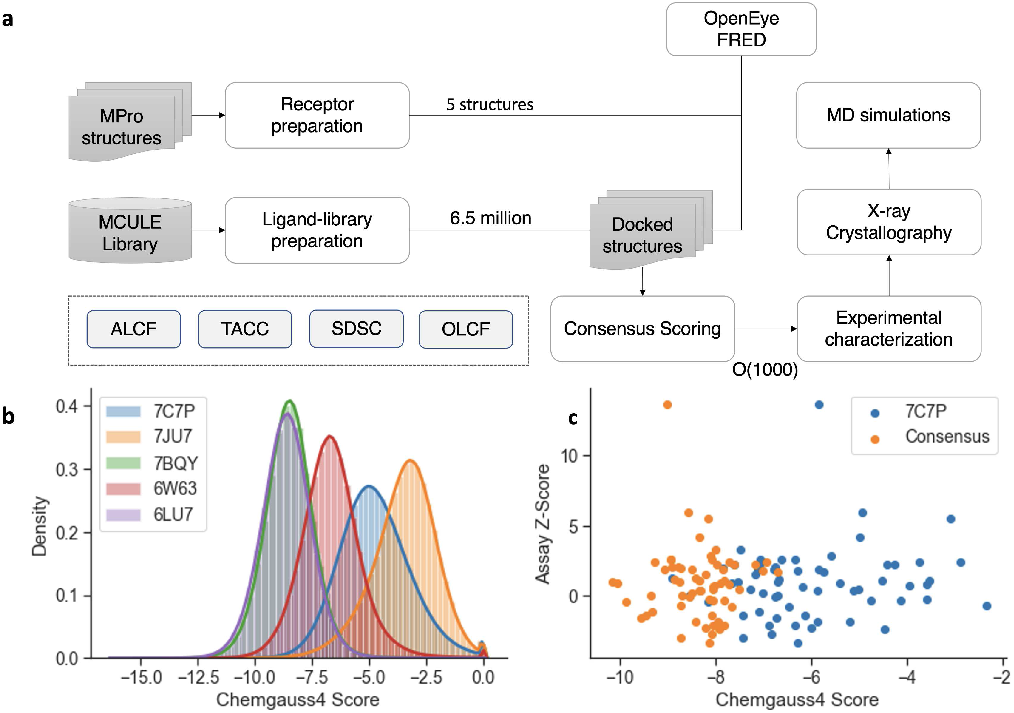
(A) Computational workflow used for screening on-demand chemical libraries against SARS-CoV2 M^pro^ with computational docking techniques. Four major supercomputing centers were utilized, namely Argonne Leadership Computing Facility (ALCF), Texas Advanced Computing Center (TACC), San Diego Supercomputing Center (SDSC), and Oak Ridge Leadership Computing Facility (OLCF). (B) The distribution of Chemgauss4 scores, from docking, from the docking a 6 million in-stock compound library. (C) The consensus scoring used shifted possible hits (higher Z-score is better) towards better scoring regions over just a single score from a single structure (7C7P is used for illustration). A lower consensus score implies a higher likelihood from the docking programs that the candidate compound will bind to the receptor.

The resulting compounds were ranked based on the docking scores in conjunction with visual inspection and availability at the time, and selected compounds were ordered for experimental validation studies. Interestingly, docking score distributions across each of the structures were slightly different (summarized in Fig. 1b-c), and we therefore examined the top 0.1% of the overall distributions. Between receptors’ respective docking, the highest correlation coefficient is 0.85 (7BQY and 6LU7) and the lowest was 0.001 (6W63 and 7JU7; See SI Table S1). In fact, 6W63’s docking result is an outlier with respect to the other four receptors, with the highest correlation coefficient of only 0.003. Given the variation amongst docking results between receptors, a consensus score was deemed necessary. A consensus score was created by taking the minimum over the available series. A minimum was chosen rather than an average, or other aggregation techniques, due to the nature of our docking protocol. OpenEye FRED has a wide range of scores, unbounded above or below. A small steric difference between receptors can cause a wide numerical discrepancy or even lack of a result. We see in Fig. 1(d) a significant difference between the correlation of consensus scores over using single samples (Table S1).

### B. MCULE-5948770040 is a SARS-CoV-2 M^pro^ inhibitor

Based on the consensus scoring procedure above, 116 compounds from the Mcule database were selected for experimental screening using the top 20 from different M^pro^ crystal structures. Of these 116 compounds, five were not available for ordering, 15 were excluded due to pan assay interference compounds (PAINS) violations based on the substructure filters of Baell and Holloway (27), and 72 were ultimately delivered. These compounds were subjected to a primary SARS-CoV-2 M^pro^ activity inhibition screen in which they were pre-incubated with the enzyme and the initial velocities of cleavage of a fluorescence resonance energy transfer (FRET) peptide were determined (28). The Z’-factor of the assay was 0.65, and the distribution of z-scores of compounds and positive (no inhibitor) and negative (no enzyme) controls is shown in Fig. 2A. At least 25% inhibition was observed for seven compounds, with MCULE-5948770040 resulting in the lowest residual activity at 20 *µ*M (12%, SI Table S2).

**Fig. 2.**
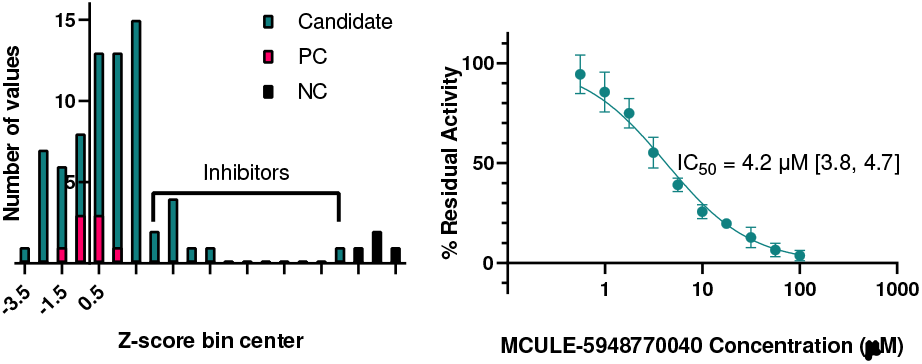
Plate-based M^pro^ activity inhibition screening and hit confirmation. (A) Histogram of z-scores of candidate inhibitors, no enzyme negative controls (NC), and no inhibitor positive controls (PC). (B) Inhibition of M^pro^ activity *in vitro* with increasing concentration of MCULE-5948770040. Initial rates are normalized to no inhibitor control (100% activity) and no enzyme control (0% activity). Error bars are standard deviation of two independent experiments, each performed in triplicate. Lines indicate the nonlinear regression of the [Inhibitor] vs. normalized response IC_50_ equation to the data with GraphPad Prism. Bracketed values indicate 95% confidence intervals from the regression.

#### Mechanism of inhibition

The concentration-dependence of MCULE-5948770040 *in-vitro* M^pro^ inhibition was measured at 40 *µ*M substrate, giving an IC_50_ of 4.2 *µ*M [95% confidence interval 3.8, 4.7] (Fig. 2B). An orthogonal quantitative high-throughput mass spectrometry-based endpoint assay was also performed at 40 *µ*M unlabelled peptide substrate, giving a similar IC_50_ of 2.6 *µ*M [95% CI 2.3, 2.9] (Fig. S2). Initial rates measured at 20-500 *µ*M substrate and 0-25 *µ*M inhibitor were consistent with a competitive mechanism of inhibition with a *K*_*i*_ of 2.9 *µ*M (Fig. S2).

### C. Room-temperature X-ray crystal structure of M^pro^ in complex with MCULE-5948770040

To elucidate the molecular basis of M^pro^ inhibition by the MCULE-5948770040 compound, a X-ray crystal structure of M^pro^ in complex with the compound was determined to 1.80 Å at near-physiological (room) temperature (Table S3 and Fig. S3). The M^pro^/MCULE-5948770040 complex crystallized as the biologically relevant homodimer with the protomers related by a two-fold crystallographic axis (Fig. 3). The tertiary fold is shown in Fig. 3. Each protomer consists of three domains (I-III). The substrate-binding cleft is formed at the interface of catalytic domains I (residues 8-101) and II (residues 102-184), whereas the *α*-helical domain III (residues 201-303) creates a dimerization interface (22, 29). The substrate binding cleft lies on the surface of the enzyme and accommodates amino acid residues or inhibitor groups at positions P1’-P5 in subsites S1′-S5, respectively (30–32). Subsites S1, S2, and S4 have well defined shapes while S1′, S3, and S5 are surface-facing with poorly defined edges (33). The non-canonical catalytic dyad composed of Cys145 and His41 lies deep within the substrate binding cleft poised for peptide bond cleavage between the C-terminal P1′and N-terminal P1 positions.

**Fig. 3.**
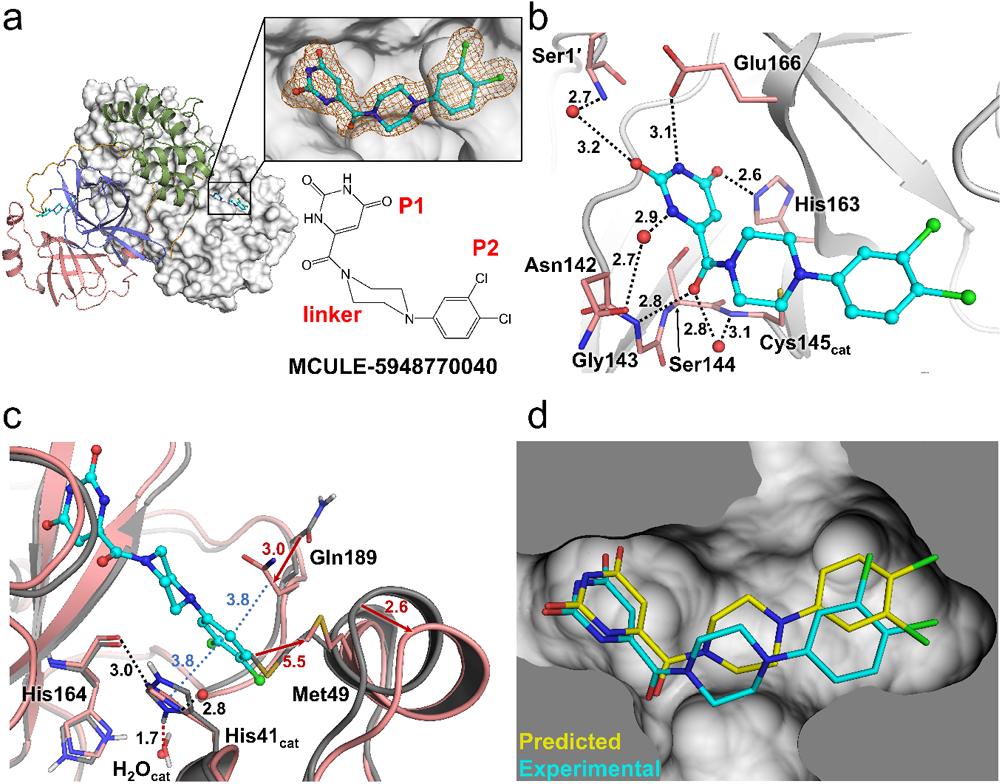
Room-temperature X-ray crystal structure of M^pro^ in complex with MCULE-5948770040 and comparison with ligand-free and docked structures. A) Overall M^pro^ homodimer in complex with MCULE-5948770040 (Cyan carbon ball and stick representation). One protomer is shown as a cartoon representation with domains I, II, and III in pink, purple, and green respectively and orange interdomain loops. The other protomer is shown as white surface. Insets show MCULE-5948770040 electron density (2Fo-Fc at 1.2σ as orange mesh) and 2D chemical diagram. B) Intermolecular interactions between M^pro^ (grey cartoon with salmon sticks) and the ligand. H-bonds are shown as black dashes. Distances in Å. C) Superposition of the M^pro^/9MCULE-5948770040 complex (salmon) with ligand-free X-ray/neutron structure (grey, PDB code 7JUN). Red arrows indicate conformational shifts from ligand-free structure to complex structure. Blue dots show π − π interactions with the P2-dichlorobenzene group. Red dashes represent a lost H-bond due to catalytic His41’s imidazole side chain flip. D) Comparison of computationally predicted (yellow carbons) and experimentally determined (cyan carbons) pose of MCULE-5948770040 bound to M^pro^.

MCULE-5948770040 binds non-covalently to the active site of M^pro^, occupying subsites S1 and S2. The electron density for the inhibitor is unambiguous (Fig. 3a) enabling accurate determination of the protein-ligand interactions (Fig. 3b). The uracil P1 group of the ligand is situated in the S1 subsite driven by polar contacts. N*ϵ*2 of the His163 imidazole side chain makes a close 2.6Å H-bond with the carbonyl at position 4 of the uracil substituent. Notably, His163 was previously determined to be singly protonated on the N*δ*1 by neutron crystallography (33), suggesting a possible rearrangement of the protonation state for His163 side chain upon ligand binding. The far end of the S1 subsite is formed such that the second protomer’s N-terminal protonated amine creates H-bonds with the Glu166 sidechain, Phe140 main chain carbonyl, and a water molecule. The amide NH at position 3 of the ligand’s P1 heterocycle is situated within hydrogen bonding distance with Glu166 and Phe140 although the geometry is unfavorable. The carbonyl and amide NH at positions 2 and 1 participate in water-mediated H-bonds with Ser1′and Asn142, respectively. M^pro^ features an oxyanion hole created by the main chain amide NH groups of Gly143, Ser144, and Cys145 at the S1 subsite base. A carbonyl linking the P1 uracil to the central piperazine linker of MCULE-5948770040 is positioned on the perimeter of the oxyanion hole forming a direct 2.8Å H-bond with Gly143 and a water-mediated contact with Cys145. Piperazine is located above the catalytic Cys145 side chain that was determined to be a deprotonated, negatively charged thiolate in the neutron structure of the ligand-free M^pro^. The P2 dichlorobenzene substituent occupies the largely hydrophobic S2 subsite.

An overlay of the MCULE-5948770040 complex with the ligand-free joint neutron/X-ray crystal structure of M^pro^ (33) shows that the P2-dichlorobenzene group moves into the hydrophobic S2 pocket altering the position of the Met49 side chain and pushing out the P2-helix (residues 46-50) by as much as ∼2.6 Å (Fig. 3c). The Met49 terminal methyl is shifted ∼5.5 Å away and its C*α* atom moves by ∼1.1 Å. Furthermore, the position of the P2-dichlorobenzene group is stabilized by the ß-ß stacking interactions with Gln189 and the imidazole side chain of catalytic His41 with the interatomic distances of ∼3.8 Å. The Gln189 side chain amide is recruited from 3 Å away from its position in the ligand-free structure and the C*α* atom shifts by almost 1 Å. Thus, the P2-dichlorobenzene is sandwiched between the side chains of these two residues. Interestingly, the binding of MCULE-5948770040 to the M^pro^ active site cleft causes the His41 side chain to flip and a *χ*2 angle rotation to create a favorable geometry for ß-ß stacking with P2-dichlorobenzene in the complex structure. Such change in the His41 conformation severs a conserved H-bond between the His41 N*δ*1 and the conserved catalytic water molecule (H_2_O_cat_) normally seen in M^pro^ structures (34, 35), but replaces it with a direct H-bond to the main chain carbonyl of His164 and results in recruitment of an additional water molecule from the bulk solvent to make an H bond with the His imidazole ring.

The computationally predicted binding pose of MCULE-5948770040 to the M^pro^ active site is in good agreement with the experimentally determined orientation (Fig. 3d). Only minor discrepancies in the piperazine linker and P2-dichlorobenzyl are present. The ligand’s uracil group forms the same polar interactions in the docked pose as observed in the crystal structure. The piperazine linker is best modeled as a chair conformation in the crystal structure while the docked geometry scored highest with it adopting a twisted boat. P2-dichlorobenzene fits well into the S2 pocket as observed in the experimental configuration, despite a 180° rotation of the aromatic ring.

### D. M^pro^interacts with MCULE-5948770040 through conformational changes within the binding site

In order to understand how the molecule interacts with the M^pro^ binding site, we carried out *µ*s-timescale atomistic molecular dynamics (MD) simulations (see Methods). For the ligand-bound (LB) state simulations, we modeled the ligand present in both M^pro^ protomers. Within the timescales of our simulations (*µ*s timescales), we observed that the ligand stays bound to the primary binding site (for both chains A and B in the dimer). The protein also does not undergo any large conformational changes (as seen from the root-mean squared deviations/ RMSD from the starting structure SI Fig. S4-S6).

We used the root mean squared fluctuations (RMSF) from ligand-free (LF) and ligand-bound (LB) M^pro^ simulations (Fig. 4A) to understand how the ligand impacts the conformational dynamics. For convenience, we considered each protomer individually (although the simulations were run with the ligand bound to both monomers in the active dimer form) and observed that several distinct regions across M^pro^ exhibit altered fluctuations. In all of our replicas, the RMSF in chain A of the dimer were slightly higher than in chain B. Distinct regions within M^pro^ respond to the ligand (rounded rectangles in Fig. 4A); these mostly consist of flexible loops surrounding the immediate vicinity of the binding pocket, corresponding to the sites (S1 - orange rounded rectangle and S2 - red rounded rectangle). Other regions surrounding the binding site (S3 – green and S4 – blue rounded rectangles respectively) also exhibit stabilization upon ligand binding. However, it is notable that not all regions exhibit stabilization within each protomer (e.g., region S4, the protomer chain A exhibits similar fluctuations to the ligand-free state). Interestingly, regions farther away from the binding site, including domain III of each protomer (R5 in Fig.4 (purple rounded rectangle) exhibit lower fluctuations in the LB simulations.

**Fig. 4.**
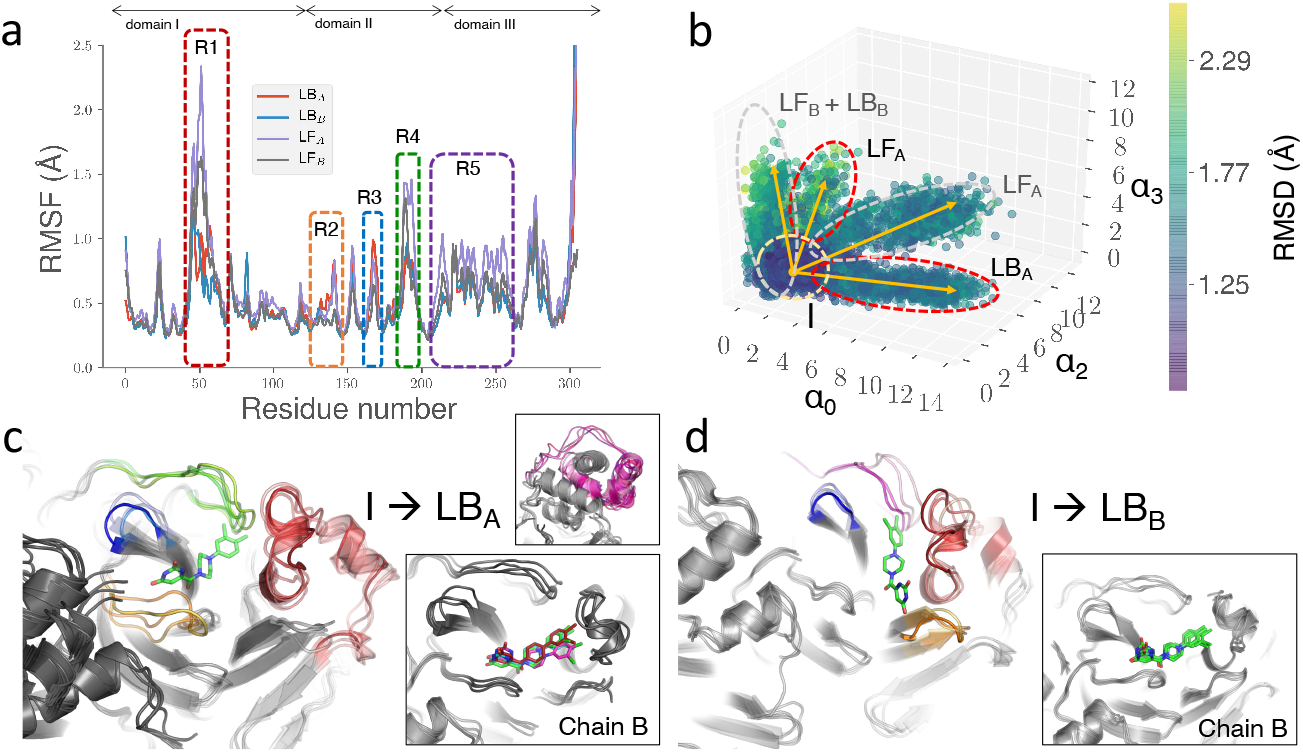
Conformational changes upon MCULE-5948770040 binding to M^pro^ indicate changes within distinct regions, both close-to and farther-away from the primary binding site. (a) RMS fluctuations of the LF- and LB-state of M^pro^ show several regions with decreased fluctuations that are highlighted within rounded rectangles. Although several regions within these regions are largely similar, amino-acid residues interacting with the ligand stabilize the binding site. (b) To further quantify the nature of these fluctuations, we characterized the collective motions which shows distinct conformational states sampled by the ligand-free (LF) and ligand-bound (LB) states. The yellow arrows indicate conformational transitions from the average structure towards the distinct conformational states (I, LF_*A*_, LF_*B*_, LB_*A*_ and LB_*B*_). These transitions are mapped in (c) I →LB_*A*_ and (d) I →LB_*B*_. (We show the I →LF_*A*_ and I →LF_*B*_). In each case, we observed that M^pro^ chain B of the dimer was more stable the chain A (insets). Regions highlighted in (a) show the motions undergone by the different regions of M^pro^.

To elucidate the collective motions that are influenced by ligand binding, we used anharmonic conformational analysis enabled autoencoder (ANCA-AE; see Methods) to embed the conformational landscape spanned by the ligand-free and ligand-bound simulations in a low-dimensional manifold (summarized in Fig. 4b). Notably, our simulations can be embedded within an ten dimensional manifold (Fig. S7) which best explains conformational fluctuations undergone by the protein. Of these embeddings, the LF and LB simulations occupied distinct projections, with the LF-simulations sampling diverse conformational states (as quantified by the RMSD to the LB states). The predominant conformational changes in the LB simulations were confined to the binding pocket spanning S1-S4 and R5, shown in Fig. 4c-d in protomer A, while we did not observe significant motions with respect to protomer B (shown in inset). The fluctuations observed were mostly a consequence of reorienting the ligand within protomer A from its primary interaction site (P1,P2) to cover the complementary binding site of (P2,P4; orange rounded rectangle in Fig. 4c, labeled I →Holo_*B*_). Notably the P1-uracil forms new interactions with the S4 region (residues) while the P2-dichlorobenzene stays bound within the hydrophobic pocket. Although we do not observe such a ligand placement within the crystal structure, these ligand motions are prevalent across multiple replicas of our simulations and can form stable interactions between the uracil and protein side-chains. Corresponding to these changes, motions in domain III of the protein (purple cartoon insets in Fig. 4 c-d) are also suppressed, showing that this region may be stabilized upon ligand binding. We also examined the hydrogen bonding patterns between the ligand and protein from the LB-simulations (SI Fig. S8) and found that the hydrogen bonds between the ligand and the protein in protomer chain B are more stable than the hydrogen bonds in chain A.

## 3. Discussion

Our study demonstrates a 2.9*µ*M potent M^pro^ inhibitor discovered through HTVS. After computationally screening 6 million molecules, we located 100 promising compounds based on a consensus score across five different M^pro^ crystal structures. Through X-ray crystallographic studies, we observed that the compound MCULE-5948770040 forms stable interactions within a hydrophobic pocket (S2) formed by the P2-dichlorobenzene group along with the P1-uracil group occupying the S1 site. Assay results also indicate that this molecule is a *µ*M non-covalent inhibitor of this enzyme and can act as a competitive inhibitor. Our *µ*s timescale simulations indicate fairly stable interactions between the protein and the ligand, which suggest that several regions of M^pro^ – both in the vicinity of the binding site and distal from it – are impacted upon binding. This alters the conformational states accessed by the protein as quantified by our ANCA-AE approach.

The combination of experimental validation with computational tools is essential to the rapid development of an inhibitor for M^pro^. Without the ability to quickly obtain a compound and experimentally validate it, computational results alone will not be a solution. Virtual screens against M^pro^ have ranged from small libraries of existing approved pharmaceuticals to natural product libraries and billion-scale combinatorial libraries (19, 36). Studies such as (37) use drug repurposing databases that are smaller (<50k) but have potential for faster lead to drug time. In this study, we aimed to balance library size and feasibility of validation, hence we opted for a 6M in-stock chemical library from Mcule. While used a approximately 120 million compound library to obtain hits for D4 dopamine receptor, we were able to use our approximately 6 million compound library without any advanced filtering post consensus scoring to locate a *µ*M hit for M^pro^.

The collective conformational motions elucidated using ANCA-AE suggest an intrinsic asymmetry in how the ligand interacts with the two protomers. Our simulations point to a mechanism of complementary interactions and inter-domain motions whereby the ligand stabilizes the conformations of the loops around the binding site as well as a loop within domain III that is considerably far away from either binding sites. In order to elucidate if the ligand binding affects either protomer separately, we also carried out simulations where the ligand was bound to only one of the two protomer units. Our analysis (SI text and Fig. S10-S11) of these simulations further indicate that the fluctuations in domain III of the protein are only affected from the ligand-bound chain. Taken together, our simulations suggest that the primary mechanism by which MCULE-5948770040 binds to and interacts with M^pro^ is by stabilizing the loops in and around the binding site. The binding of the ligand is asymmetric in the protomers; while it is stable in one of the protomers, it undergoes a slight conformational change (albeit stable) within the binding pocket while still maintaining the strong hydrophobic interactions within P2. Further, our analysis indicates that the hydrogen bonding patterns are different for the two chains, which also lends support to the idea that the collective motions as induced by the LB-states may indeed be different.

Computational protocols for HTVS vary widely across existing SARS-CoV-2 M^pro^ screens, from accurate but expensive simulations for MMGBSA/PBSA scoring to standard docking. Gorgulla et al. (19) screened Enamine Real, a billion-scale combinatorial product library, against various SARS-CoV-2 targets using QuickVina W—a slightly less accurate but computationally efficient flavor of AutoDock Vina (39). Acharya et al. (13) also screened Enamine Real with Autodock-GPU. In comparison one of the few other studies which involve assay and crystallographic experimental studies for M^pro^ lead generation (17), our work relies on a single computational workflow rather than community lead sourcing (where the methods of each contributor are not restricted). Rather than taking community input for prioritizing experimental leads, our work features an HTVS protocol based on ensemble docking with consensus scoring. Using the web portal to access hits from (17), we compared the difference between leads from domain experts with the library we used for screening and found that both groups arrived at structurally similar hits independently, with the same P1-Liner-P2 topology and interaction with M^pro^ S1 and S2 sites (SI Fig. S12). Between the two groups, our best compounds share piperazine and the uracil groups.

The compound MCULE-5948770040 forms stable interactions with both the protomers as we observed from our X-ray crystallography and MD simulations. Our simulations provide insights into how the conformational fluctutations of the protein are altered in response to the ligand binding to the primary site. In fact, we observed that the fluctuations in region R5, which is over 20 Å away from the primary binding site in M^pro^, are effected by the ligand binding. Further, the ligand’s interaction with M^pro^ alters the conformational states accessed by the enzyme, notably along the substrate binding loops. Compared to other ligands that have been structurally characterized (as well as the substrate peptide), MCULE-5948770040 is much smaller and interacts stably with both the S1 and S2 sites within M^pro^. Thus, a design strategy that targets the S1-S2 sites and mimics important features of the peptide side chains is sufficient for identifying inhibitors of M^pro^. We have early indications that molecules structurally similar to MCULE-5948770040 also demonstrate inhibition, and efforts are currently underway to identify promising candidates, which we expect to publish shortly.

## Materials and Methods

### A. Molecular library generation

We use a set of on demand compounds from Mcule (ORD), which can be freely obtained on their website (16). ORD consists of compounds from Mcule listed as available on their website, Mcule Purchasable (Known Stock Amounts).

### B. Molecular Docking Protocol with OpenEye Toolkit

The main protease structures screened included the following PDB structures: 7BQY, 6LU7, 6W63, 7C7P, and 7JU7. The receptors for OpenEye Chemgauss4 scoring were prepared using the known binding region of M^pro^ with the OpenEye Docking Toolkit(20)

For memory efficiency, conformer generation and tautomerization were performed on-the-fly. OMEGA (40) was used, sampling around 300–500 conformations for each ligand. When ligand-binding information was available in the receptor, HYBRID was used due to its increased pose prediction accuracy over FRED (20). HYBRID and FRED have the same scoring function; however, HYBRID uses a heuristic to reduce the search space for ligand positioning. The best score from the ensemble of tautomers and conformers is chosen as the representative “docking score” for the chemical species.

### C. Computational Workflow

OpenEye toolkit’s FRED docking program was deployed on Frontera at TACC. Docking scores for the M^pro^ receptor were computed with individual runs per pocket. The docking protocol as described above requires the following steps for each compound: (1) load receptor into memory; (2) load compound data stored in MCULE (16) database from disk; (3) run the specific docking protocol over the receptor/compound pair; and (4) write the resulting docking score to persistent storage.

High-throughput docking was implemented using RADICAL-Pilot (RP) and RAPTOR (41). RP is a pilot-enabled runtime system while RAPTOR is a scalable master/worker overlay developed to improve the execution performance of many, short-running tasks encoded as Python functions. The runs used between 128 and 7000 concurrent nodes. For each run, we measured throughput (the number of docking calls per hour) and resource utilization (the fraction of time that acquired nodes were kept busy). Resource utilization was dependent on the size of the run (number of compounds to dock, number of nodes to use), and was typically above 90%. In Table 1, we summarize the results from multiple runs, including one on 7000 nodes.

**Table 1.** **Resource Utilization when docking 126 × 10^6^ and 205 × 10^6^ ligands with OpenEye using RAPTOR on Frontera. Docking time varies depending on the physical properties of each ligand, affecting the obtained docking throughput.**

| #Nodes | #Ligands<br>[$\times 10^6$] | Utilization<br>[%] | Docking Time [sec] | | | Throughput [$\times 10^6$ docks/hr] | |
| --- | --- | --- | --- | --- | --- | --- | --- |
|  |  |  | min | max | mean | max | mean |
| 128 | 205 | 89.6 | 0.1 | 3582.6 | 28.8 | 17.4 | 5.0 |
| 3850 | 126 | 95.5 | 0.1 | 833.1 | 25.1 | 27.5 | 19.1 |
| 7000 | 126 | 90.0 | 1.0 | 180.0 | 10.1 | 144.0 | 126.0 |

The runtimes for individual docking calculations have a long-tail distribution (see Table 1) The mean docking time determines the achievable maximal throughput For example, the last row in Table 1 suggests an upper limit on average docking call throughput of 139M docking calls per hour (1hr * cores per node * number of nodes / mean docking time seconds = 3600 * 56 * 7000 / 10.1 = 139,722,772 ∼ 139M docks/hr). The achieved average throughput was ∼126M docks/hr.

### D. SARS-CoV-2 M^pro^ expression and purification

A gene construct encoding M^pro^ (NSP5) from SARS-CoV-2 was cloned into plasmid pD451-SR (Atum, Newark, CA), which was developed in (11), and expressed and purified consistent with the protocols detailed in (42). Protein purification supplies were purchased from Cytiva (Piscataway, New Jersey, USA). Briefly, authentic N-terminus is achieved by a NSP4-NSP5 autoprocessing sequence (SAVLQ↓SGFRK where the arrow indicates the scissile bond) flanked by maltose binding protein and M^pro^. Following M^pro^, a sequence encoding the human rhinovirus 3C (HRV-3C) cleavage site (SGVTFQ↓GP) is followed by a His6-tag. The N-terminal sequence is created by autocleavage during expression while the C-terminus is generated by HRV-3C treatment following Ni immobilized metal affinity chromatography.

### E. Primary M^pro^ inhibition screen

Compounds were purchased from Mcule, Inc. as 10 mM stock solutions in DMSO and stored at −20 °C. The assays were performed in 40 *µ*L total volume in black half area 96-well plates (Greiner PN 675076) at 25 °C. The assay buffer contained 20 mM Tris-HCl pH 7.3, 100 mM NaCl, 1 mM EDTA, and 2 mM reduced glutathione (6.15 mg added per 10 mL buffer fresh for each experiment) with 5% v/v final DMSO concentration. M^pro^ initial rates were measured using a previously established fluorescence resonance energy transfer (FRET) peptide substrate assay(28). The FRET substrate DABCYL-KTSAVLQ↓SGFRKM-E(EDANS) trifluoroacetate salt was purchased from Bachem (PN 4045664), dissolved to 10 mM in DMSO and stored in aliquots at −20 °C. 10 *µ*L enzyme solution was dispensed into wells (250 nM final concentration), followed by 10 *µ*L inhibitor solution (20 *µ*M final concentration), centrifuged briefly, and incubated for 30 min. Reactions were initiated by adding 20 *µ*L substrate at 40 *µ*M final concentration. Fluorescence was detected every 24 s by a Biotek Synergy H1 plate reader with an excitation wavelength of 336 nm and an emission wavelength of 490 nm, 6.25 mm read height, low lamp energy, and 3 measurements per data point. After background subtraction of the average of no enzyme negative controls, product formation was quantified using a 0.05 – 22 *µ*M calibration curve of the free EDANS acid (Sigma PN A6517). Product concentrations were adjusted for inner filter absorbance effects with correction factors generated by comparing the fluorescence of 2 *µ*M EDANS in solution with each concentration of substrate used to that with no substrate. Initial rates were determined for time points in the linear range by linear regression in Excel, residual activities were determined by normalizing candidate initial rates to the average of the positive controls, and z-scores were determined by dividing the difference between the candidate initial rate and average positive control initial rate by the standard deviation of the positive control initial rates. The Z’ statistic for the plate was calculated using the published equation.

### F. Peptide synthesis

The unlabelled M^pro^ substrate peptide AVLQ↓SGFRKK-amide and the isotopically labelled substrate and product peptide internal standards (A+7)VLQSGFRKK-amide and (A+7)VLQ-OH were synthesized by automated peptide synthesis using a Liberty PRIME™ peptide synthesizer (CEM). Reagents were peptide synthesis or biotechnology grade. Amino acids were purchased from P3Bio, and Fmoc-[^13^C_3_, ^15^N, D_3_]-alanine (A+7) was previously synthesized at Los Alamos National Laboratory following published protocols (43). Other purchased reagents were dimethylformamide (DMF), pyrrole (prepared as 20% v/v in DMF), and high performance liquid chromatography (HPLC)-grade acetonitrile (Alfa Aesar), diisopropylcarbodiimide and Oxyma Pure (AKScientific), N,N-diisopropyl ethyl amine (DIPEA), triisopropyl silane (TIPS), trifluoroacetic acid (TFA), thioanisole, Rink amide resin, and octaethylenglycol-dithiol (Sigma Aldrich), dichloromethane and Optima mass spectrometry grade acetonitrile (Fisher Scientific).

Peptide syntheses were performed at 0.1 mM scale under argon on a Rink amide resin with 0.1 M DIPEA added to the Oxyma solution to prevent hydrolysis of acid labile side chain protecting groups, obtaining average yields for double couple cycles of >99%. For stable isotope labeled peptides only two equivalents of the labeled amino acid were used and coupling time was extended to 20 min at 90 °C. Peptides were deprotected with the following mixture: 1.25 mL TIPS, 0.625 mL thioanisole, 1.25 mL octaethyleneglycodithiol, and after 5 min TFA was added to a total volume of 25 mL. Solutions were filtered and the filtrate concentrated to 10 mL, followed by precipitation with ice cold ether and collection by centrifugation.

Peptides were purified to >98% by Waters HPLC workstation (2545 pump with 2998 photodiode array detector) with a Waters BEH 130, 5 *µ*m, 19×150 mm C18 column and a linear gradient from 98:2 to 50:50 water:acetonitrile with 0.1% TFA at 20 mL/min. Absorbance at 215 nm was monitored and peaks were collected and lyophilized to yield a white fluffy solid. Peptide purity was analyzed by analytical HPLC and Thermo LTQ mass spectrometry with electrospray ionization in positive mode with a Waters BEH 130, 5 *µ*m, 4.6×150 mm C18 column and a linear gradient from 96:2 to 60:40 water:acetonitrile with 0.1% TFA at 1.5 mL/min over 12 min (Fig. S2).

### G. Quantitative mass spectrometry M^pro^ inhibition assay

The quantitative mass spectrometry (MS) inhibition assay was performed as described for the FRET-based primary screen with some modifications. Round-bottom polypropylene 96-well plates (Corning PN 3365) were used with 150 nM final M^pro^ concentration and the unlabeled peptide substrate synthesized above. Five min after substrate addition, the assay was quenched 1:1 v:v with 2% formic acid in water with 2 *µ*M of each internal standard peptide from above, centrifuged 10 min at 4 °C, and the supernatant was diluted 1:9 v:v into 1% formic acid. Substrate and product peptides and internal standards were quantified by high-throughput MS using a Sciex 5500 QTRAP with a custom open port sampling interface (OPSI) (44). Samples were introduced as 2 *µ*L droplets and the OPSI-MS analysis was performed using 10:90:0.1 v:v:v water:methanol:formic acid at 80 *µ*L/min. Positive ion mode electrospray ionization parameters were CUR: 25, IS: 5000, TEM: 400, GS1: 90, GS2: 60, EP: 10, and CXP: 10. Optimized multiple reaction monitoring detection parameters were dwell: 50 msec, product DP: 100 and CE: 25, and substrate DP: 150 and CE: 34. The following mass-to-charge transitions were monitored: substrate AVLQSGFRKK, 566.9→722.3; (A+7)VLQSGFRKK, 570.4→722.3; product AVLQ, 430.3→260.3, and (A+7)VLQ, 437.3→260.3. Product formation and remaining substrate were quantified by dividing the peak area of the transitions by that of the corresponding internal standard transitions.

### H. IC_50_ and *K*_*i*_ Value Determination

To determine the concentration at which a compound was able to achieve 50% inhibition of M^pro^ activity *in vitro* (IC_50_), the FRET and quantitative MS assays described above were performed at 10 concentrations of inhibitor (0.56-100 *µ*M) in triplicate with 150 nM enzyme. Initial rates, for FRET, or product formation in 5 min, for MS, were normalized to no inhibitor control (100% activity) and no enzyme control (0% activity), and nonlinear regression of the [Inhibitor] vs. normalized response IC_50_ equation was performed to fit the data using GraphPad Prism 9.0.0, yielding IC_50_ and its 95% confidence interval. To confirm the mechanism of inhibition and determine *K*_*i*_, the FRET activity assay was performed at 8 concentrations of substrate (20-500 *µ*M) and 4 concentrations of inhibitor (0-25 *µ*M) in triplicate in two independent experiments. A global nonlinear regression was performed to fit the competitive inhibition equation to the entire data set using GraphPad Prism 9.0, yielding *K*_*M*_, *K*_*i*_, *V*_*max*_, and their associated 95% confidence intervals.

### I. Crystallization

Crystallization reagents were purchased from Hampton Research (Aliso Viejo, California, USA). Crystallographic tools were purchased from MiTeGen (Ithaca, New York, USA) and Vitrocom (Mountain Lakes, New Jersey, USA). M^pro^ was concentrated to ∼5.0 mg/mL in 20 mM Tris pH 8.0, 150 mM NaCl, 1 mM TCEP, for crystallization. The presence of reducing agent such as TCEP is essential for preventing oxidation of the catalytic cysteine sidechain (45). Conditions for growing crystalline aggregates of ligand-free M^pro^ were identified by high-throughput screen at the Hauptman-Woodward Research Institute (46) and reproduced locally using 22% PEG3350, 0.1 M Bis-Tris pH 6.5 in 20*µ*L drops with 1:1 ratio of the protein:well solution using sitting-drop vapor diffusion with microbridges. Crystal aggregates of ligand-free sample were converted to microseeds with Hampton Research Seed Beads™ and used for nucleating M^pro^ crystals in subsequent co-crystallization experiments. Lyophilized MCULE-5948770040 for co-crystallization was dissolved in 100% DMSO as a 50 mM stock stored at −20°C. MCULE-5948770040 was mixed with M^pro^ at 5:1 M ratio and allowed to incubate on ice for a minimum one hour. Crystals were grown in a 40 *µ*L drop at a 1:1 mixture with 18% PEG3350, 0.1 M Bis-Tris pH 7.0 with 0.2 *µ*L of 1:200 dilution microseeds and incubated at 14°C. A large crystal measuring ∼ 1 × 0.5 × 0.3 mm suitable for room-temperature X-ray diffraction grew after 2 weeks (Fig. S2).

### J. Room-temperature X-ray data collection and structure refinement

The protein crystal was mounted using a MiTeGen (Ithaca, NY) room-temperature capillary system (Fig. S2. X-rays for crystallography were generated from a Rigaku HighFlux HomeLab employing a MicroMax-007 HF X-ray generator and Osmic VariMax optics allowing diffraction images to be collected using an Eiger R 4M hybrid photon counting detector. Diffraction data was reduced and scaled using Rigaku CrysAlis Pro software package. Molecular replacement was performed using the ligand-free room-temperature M^pro^ structure (PDB code 6WQF) (11) using Molrep (47). Structure refinement was performed with Phenix.refine from Phenix suite (48) and COOT (49) for manual refinement and Molprobity (50). Data collection and refinement statistics are listed in Table S1. The structure and corresponding structure factors of the room temperature Mpro/MCULE-5948770040 complex have been deposited into the Protein Data Bank with the PDB accession code 7TLJ.

### K. Molecular dynamics simulations of M^pro^ complex with MCULE-5948770040

The crystal structure of protein dimer was modeled with the AMBER molecular modeling package (51) with the amber.ff14sb force field parameters (for the protein) (52) and with the GAFF parameters (for the ligand) (53). In order to better determine the partial charges for the ligand, quantum mechanical (QM) calculations were performed using NWChem (54) based on the RESP method at B3LYP/6-31G* level of theory (55), while all bonded parameters were taken from GAFF force field.

The systems (both the ligand-bound/LB and ligand-free/LF) were solvated using the TIP3P water model and counter ions were added to neutralize the charge. After equilibrating the systems using previously published protocols (56), we carried out production runs using the OpenMM (57) simulation package on Nvidia V100 GPUs using the Argonne Leadership Computing Facility’s (ALCF) computing clusters. Each time step was integrated with Langevin integrator at 310 K, 1 ps^−1^ friction coefficient, and 2 fs interval with fixed lengths being maintained for atomic bonds involving hydrogen atoms. System pressure was maintained at 1 atm with the MonteCarloBarostat. Nonbonded interactions were cut off at 1.0 nm and Particle Mesh Ewald (PME) was implemented for long-range interaction. The simulations were run for 1 *µ*s and 50 ps reporting interval (4 replicas).

### L. Quantifying conformational transitions in the ligand-bound and ligand-free states of MPro with Anharmonic Conformational Analysis driven Autoencoders (ANCA-AE)

Conformational fluctuations within bio-molecular simulations (and specifically proteins) show significant higher order moments (see Fig. S12); these fluctuations may relate to protein function (56, 58). To quantify such anharmonic fluctuations within our simulations, we used fourth-order statistics to describe atomistic fluctuations and to characterize the internal motions using a small number of anharmonic modes (59). We projected the original data (306 C^*α*^ atoms per chain - (*x, y, z*) coordinates) onto a 40 (or 50) dimensional space, depending on the set of simulations considered. Notably, for the ligand bound states (in both protomers), 40 dimensions covered about 95% of overall variance whereas we required 50 dimensions to cover 95% of the variance when we included the ligand bound states from just one protomer.

Given the significant non-linearity in the atomic fluctuations, we used an autoencoder to further delineate the intrinsic structure in the low-dimensional anharmonic space. Similar to approaches that use variational approximations to model molecular kinetics from MD simulations (60), we used an autoencoder architecture consisting of a symmetric encoder and decoder network. The network is composed of a single dense layer with 32 dimensions, and an 8 dimensional latent space. We trained the network for 50 epochs using the RMSprop optimizer to minimize the mean-squared error (MSE) reconstruction loss with a learning rate of 0.001, weight decay of 0.00001, and a batch size of 64. We used ReLU activation in all places except the final reconstruction layer, where we used Tanh activation. A mixture of Gaussian (MoG) model was used to cluster the conformations in the low dimensional landscape, similar to the approach outlined in (61).

## Author contributions

AC designed and implemented the virtual screening approach. SG designed, performed, and analyzed plate-based activity screens. DK purified enzyme for activity screens and crystallization, and collected diffraction data, which AK supervised. JS, VK, LC assisted with ordering compounds and performing experimental methods. HM prepared and ran simulations, which AR supervised, and AB and AT, and SC assisted with. YB, BB, TB, KC, RC, LT, MT, AT, MT, and HV assisted with scaling, designing, and running workflows across supercomputing centers, which SJ and IF supervised. MH and RS oversaw the project. All authors contributed to the manuscript.

## ACKNOWLEDGMENTS

Research was supported by the DOE Office of Science through the National Virtual Biotechnology Laboratory, a consortium of DOE national laboratories focused on response to COVID-19, with funding provided by the Coronavirus CARES Act and as part of the CANDLE project by the DOE-Exascale Computing Project (17-SC-20-SC). Anda Trifan acknowledges support from the DOE-Computational Sciences Graduate Fellowship (DOE-CSGF) under grant number: DE-SC0019323.This research used resources at the Spallation Neutron Source and the High Flux Isotope Reactor, which are DOE Office of Science User Facilities operated by the Oak Ridge National Laboratory. The Office of Biological and Environmental Research supported research at ORNL’s Center for Structural Molecular Biology (CSMB), a DOE Office of Science User Facility. ORNL is managed by UT-Battelle LLC for DOE’s Office of Science. This research used resources of the Argonne Leadership Computing Facility, which is a DOE Office of Science User Facility supported under contract DE-AC02-06CH11357. This research also used resources of the Oak Ridge Leadership Computing Facility, which is a DOE Office of Science User Facility supported under Contract DE-AC05-00OR22725. The authors acknowledge the Texas Advanced Computing Center (TACC) at The University of Texas at Austin for providing HPC resources that have contributed to the research results reported within this paper.

## Supplemental Information

**Fig. S1.**
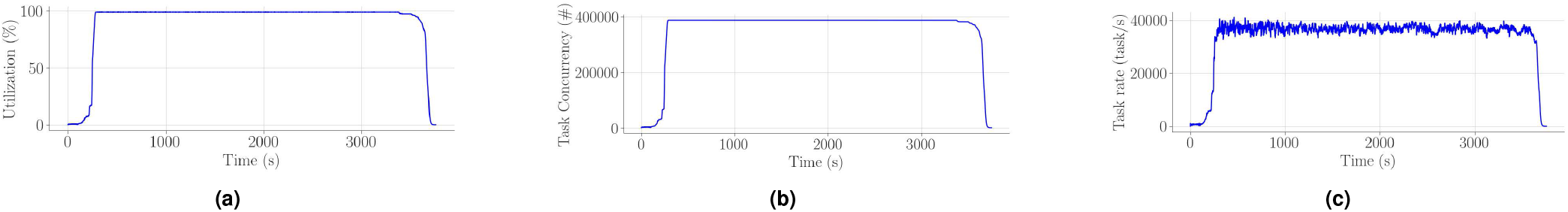
(a) resource utilization (RU), (b) execution concurrency (EC) and (c) task execution rate (TR) with RAPTOR when executing 126,471,524 OpenEye Python function calls on 7000 compute nodes/392,000 cores of Frontera with 70 master and 99 workers per master. RU = 90%; EC = 4 × 10^5^ steady state; TR = 144 × 10^6^*/hour* peak.

**Table S1.** Correlation matrix for docking scores across different receptors used.

|  | 7C7P | 7JU7 | 7BQY | 6W63 | 6LU7 | Consensus |
| --- | --- | --- | --- | --- | --- | --- |
| 7C7P | 1.000 | 0.041 | 0.395 | 0.269 | 0.400 | 0.647 |
| 7JU7 | 0.041 | 1.000 | 0.260 | 0.188 | 0.268 | 0.527 |
| 7BQY | 0.395 | 0.260 | 1.000 | 0.402 | 0.843 | 0.799 |
| 6W63 | 0.269 | 0.188 | 0.402 | 1.000 | 0.408 | 0.666 |
| 6LU7 | 0.400 | 0.268 | 0.843 | 0.408 | 1.000 | 0.806 |
| Consensus | 0.647 | 0.527 | 0.799 | 0.666 | 0.806 | 1.000 |

### 1. High Throughput Virtual Screening Workflow

To minimize the overhead caused by repeated loading of the receptor data from memory, the data were loaded once per node and then reused for all docking runs assigned to that specific node. The individual cores hosting the docking computations received cloned copies of the receptor data so as to isolate the individual docking computations. To reduce the overhead of loading compound data from disk, the storage offsets in the dataset were precomputed at startup, and shared with all the nodes, which reduced the I/O operations on the compound data. Intermediate data were stored on the local storage of each node, further reducing the load on the shared file system. For the same reason, we also stored the Python virtualenv with the OpenEye docking modules on the local storage of the nodes.

Fig. S1 shows core utilization (Fig. S1(a)), docking call concurrency (Fig. S1(b)) and docking call rate over time (Fig. S1(c)). The max docking time dominates the inefficiencies when the workload tapers off, as all idling nodes have to wait for a few remaining long running docking calls to terminate. The plots also show inefficiencies at start up, caused by: distributing input data, Python modules, and docking calls across the compute nodes; and preparing data structures in memory. Those inefficiencies lead to an average resource utilization of 91% and an average docking call throughput of 139M * 0.91 = 126M docks/hr. The achieved peak rate of 144M docks/hr indicates an uneven distribution of docking call times throughout the data set, but also shows that resource utilization during steady state is near perfect, as confirmed by Fig. S1(a).

### 2. Performance of Computational Models

Several recent papers have reported impressive performance and scaling results. However, given the diverse computing platforms and docking programs employed, as well as different measures of performance, it is difficult to provide a head-to-head comparison. We discuss two “state-of-the-art” publications, that represent the broad spectrum of performance considerations and design points.

VirtualFlow (1) submits multiple different “jobs” to different “clusters” and report a peak utilization of 160k core. However they do not report how effectively the resources are utilized and focus on application performance measures (docks/time). In contrast to our solution which submits one single job to the supercomputer and manages the entire workload within the boundaries of that single job. Our workflow on Frontera uses more than 400k cores at peak, while reporting a resource utilization consistently above 95programs are different, a direct throughput comparison is not meaningful.

In a recent submission (2) a team from ORNL demonstrated a performance of 50M docks/hour for up to 20 poses per dock (i.e., 20k docks/second) on the Summit supercomputer. This was primarily achieved through adaptation of AutoDock-GPU (3) — GPU offloading feature calculations in rescoring, and the use of GPU-accelerated database query software. Additionally, a further 10x performance improvement was achieved by using parallel database methods. In contrast, our work reports a general-purpose execution tool that is not constrained to a specific docking program, nor is it limited optimizations on a specific computing platform.

### 3. Assay details

**Fig. S2.**
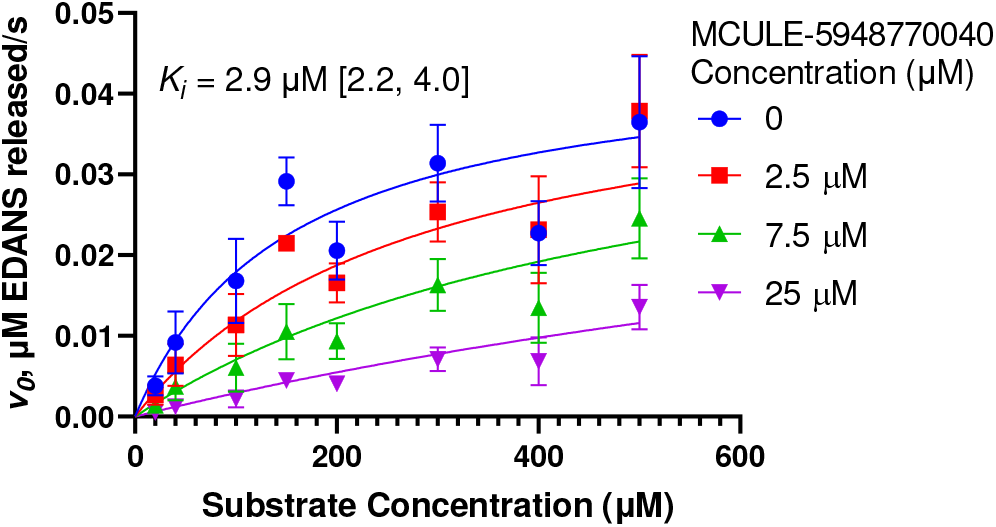
Quantitative high-throughput mass spectrometry-based endpoint assay depicting the K_*i*_ determined for MCULE-5948770040 as it binds to its substrate (M^pro^).

**Table S3.** Extended Plate-based M^pro^ activity inhibition screen results.

| Compound | Z-score | % Residual activity at 20 $\mu\text{M}$ |
| --- | --- | --- |
| PC | -0.1 | 101% |
| PC | -1.7 | 111% |
| PC | 0.8 | 95% |
| PC | -1.0 | 107% |
| PC | 1.3 | 91% |
| PC | 0.8 | 95% |
| PC | -0.1 | 101% |
| PC | 0.0 | 100% |
| NC | 16.8 | -8% |
| NC | 14.9 | 4% |
| NC | 15.4 | 1% |
| NC | 15.1 | 3% |
| MCULE-5948770040 | 13.7 | 12% |
| MCULE-9466928660 | 6.0 | 62% |
| MCULE-8841098920 | 5.5 | 64% |
| S-613265359 | 4.9 | 69% |
| S-613265357 | 4.6 | 70% |
| S-613265358 | 4.6 | 71% |
| MCULE-9437816791 | 4.2 | 73% |
| MCULE-9635688656 | 3.3 | 78% |
| S-613265352 | 3.2 | 79% |
| MCULE-5271818360 | 2.9 | 81% |
| S-613265354 | 2.9 | 81% |
| MCULE-2322094178 | 2.6 | 83% |
| MCULE-5715268546 | 2.6 | 83% |
| MCULE-6499664919 | 2.6 | 83% |
| MCULE-5087943183 | 2.4 | 84% |
| MCULE-5517230918 | 2.4 | 84% |
| MCULE-6069061169 | 2.4 | 85% |
| MCULE-9796670140 | 2.2 | 86% |
| MCULE-3840831569 | 2.2 | 86% |
| MCULE-1316421825 | 2.2 | 86% |
| MCULE-1691783948 | 2.1 | 86% |
| MCULE-9073095150 | 2.1 | 87% |
| MCULE-6576711707 | 2.0 | 87% |
| S-613265355 | 2.0 | 87% |
| MCULE-9274314730 | 1.9 | 88% |
| MCULE-6419793505 | 1.9 | 88% |
| MCULE-7344200039 | 1.9 | 88% |
| MCULE-2015327629 | 1.8 | 88% |
| MCULE-2233915378 | 1.7 | 89% |
| MCULE-5637537904 | 1.5 | 91% |
| MCULE-9408357356 | 1.3 | 92% |
| MCULE-7367467944 | 1.2 | 92% |
| MCULE-3860138249 | 1.1 | 93% |
| MCULE-4793474277 | 1.1 | 93% |
| MCULE-3980688822 | 1.0 | 93% |
| MCULE-4125658730 | 1.0 | 94% |
| S-613265351 | 1.0 | 94% |
| MCULE-8462272476 | 0.9 | 94% |
| MCULE-2988500623 | 0.9 | 94% |
| MCULE-8798249417 | 0.8 | 95% |
| MCULE-7522626224 | 0.7 | 96% |
| MCULE-2310276978 | 0.6 | 96% |
| MCULE-6998567671 | 0.5 | 96% |
| MCULE-1618528086 | 0.5 | 97% |
| MCULE-1541843003 | 0.4 | 97% |
| MCULE-5145994619 | 0.4 | 98% |
| MCULE-6929749068 | 0.2 | 99% |
| S-613265356 | 0.0 | 100% |
| MCULE-7152686229 | 0.0 | 100% |
| MCULE-8568450548 | 0.0 | 100% |
| MCULE-4811796101 | -0.1 | 101% |
| MCULE-2017284421 | -0.2 | 101% |
| MCULE-7679599807 | -0.3 | 102% |
| MCULE-5985443175 | -0.3 | 102% |
| MCULE-3102049132 | -0.4 | 102% |
| S-613265350 | -0.4 | 103% |
| MCULE-9172580823 | -0.7 | 105% |
| MCULE-9221479095 | -0.8 | 105% |
| MCULE-4489145056 | -1.1 | 107% |
| MCULE-7943829824 | -1.4 | 109% |
| MCULE-6144665861 | -1.4 | 109% |
| MCULE-1569445415 | -1.6 | 110% |
| MCULE-1224951145 | -1.8 | 111% |
| MCULE-5592055088 | -1.8 | 111% |
| MCULE-1189442017 | -2.1 | 113% |
| S-613265353 | -2.1 | 114% |
| MCULE-7291272788 | -2.1 | 114% |
| MCULE-6426577239 | -2.3 | 115% |
| MCULE-4563504276 | -2.4 | 116% |
| MCULE-6537132306 | -2.7 | 118% |
| MCULE-3593629191 | -3.0 | 119% |
| MCULE-3076850808 | -3.3 | 121% |

**Table S2.**
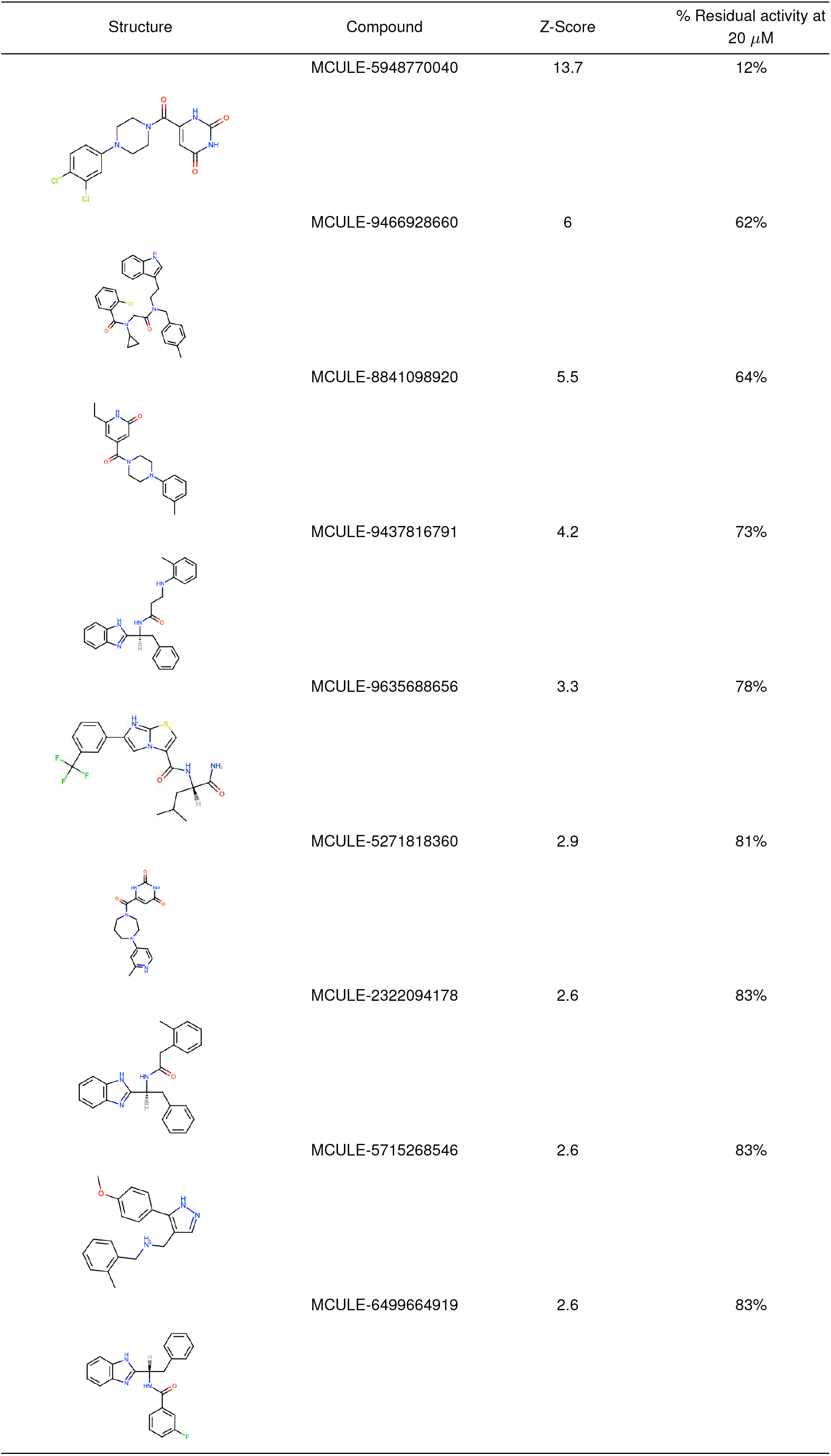
Plate-based M^pro^ activity inhibition screen results.

| Structure | Compound | Z-Score | % Residual activity at 20 μM |
| --- | --- | --- | --- |
|  | MCULE-5948770040 | 13.7 | 12% |
|  | MCULE-9466928660 | 6 | 62% |
|  | MCULE-8841098920 | 5.5 | 64% |
|  | MCULE-9437816791 | 4.2 | 73% |
|  | MCULE-9635688656 | 3.3 | 78% |
|  | MCULE-5271818360 | 2.9 | 81% |
|  | MCULE-2322094178 | 2.6 | 83% |
|  | MCULE-5715268546 | 2.6 | 83% |
|  | MCULE-6499664919 | 2.6 | 83% |

### 4. Crystal

**Fig. S3.**
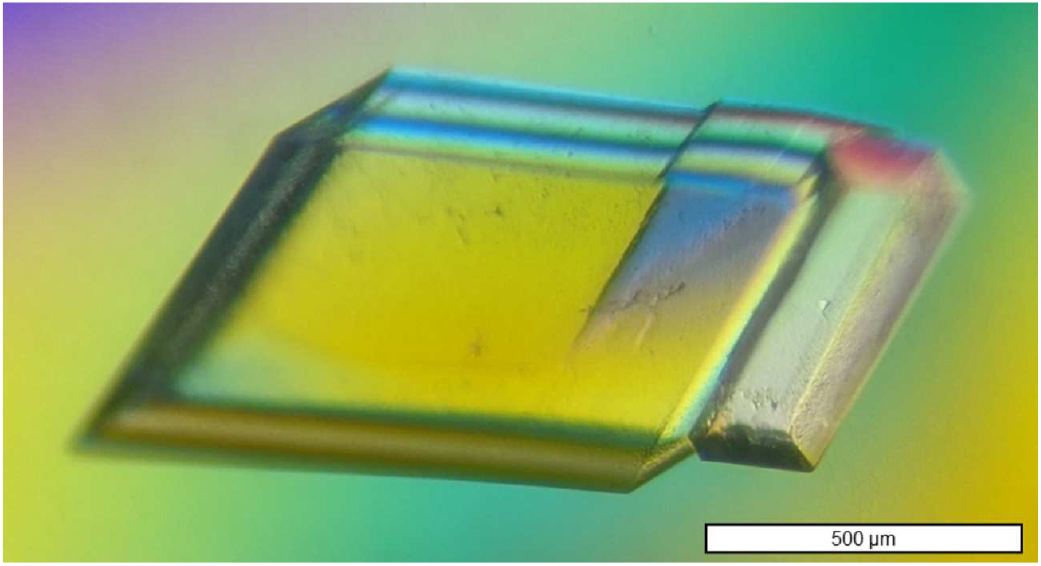
Protein crystal of SARS-CoV-2 3CL M^pro^ in complex with MCULE-5948770040 used for room-temperature X-ray data collection. Crystal measured ∼1×0.5×0.3 mm or ∼0.15 mm^3^ and diffracted to 1.80 Å using a home source X-ray diffractometer

**Table S4.**
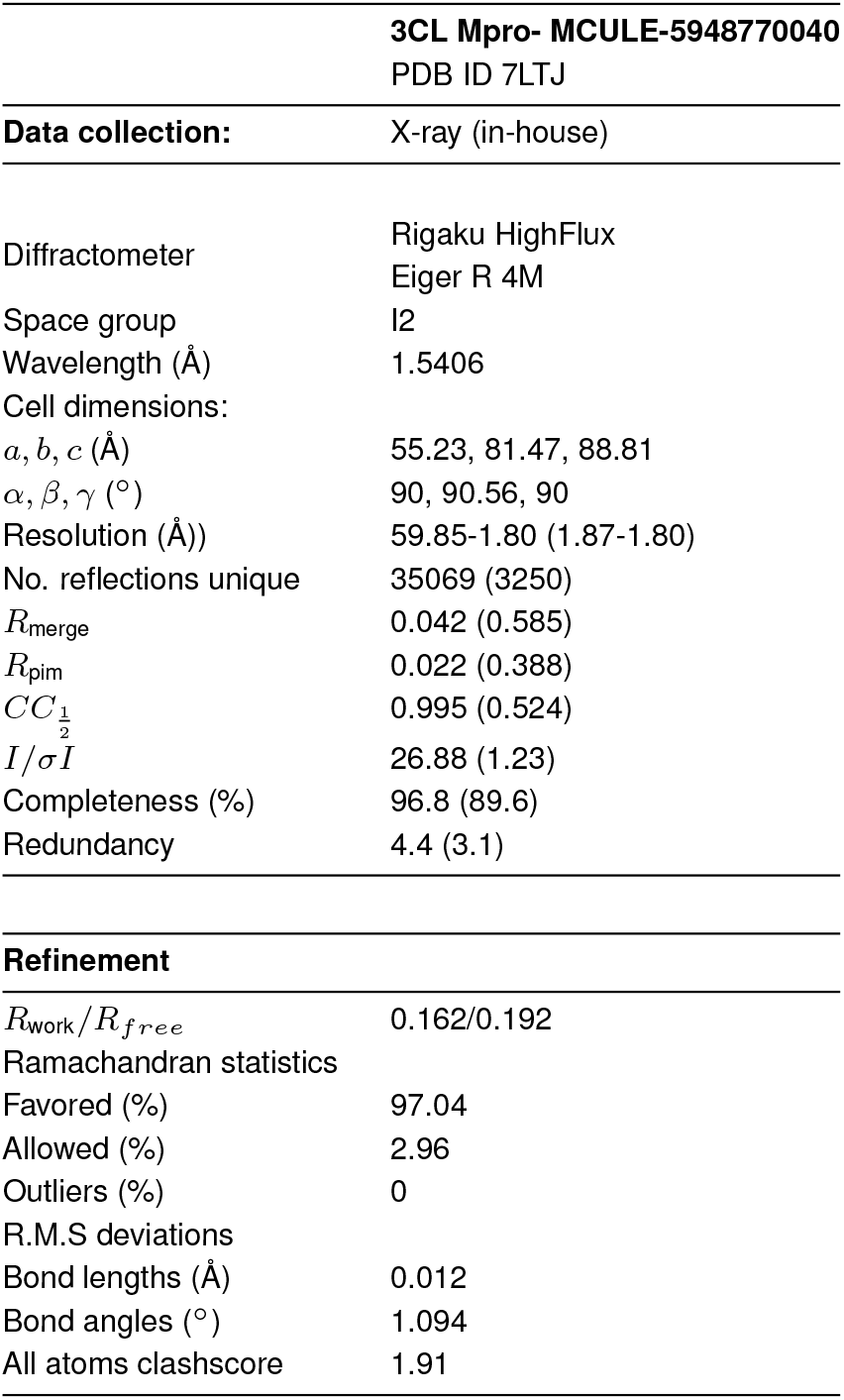
**Crystallography data reduction and refinement statistics for room temperature structure of SARS-CoV-2 3CL Mpro in complex with MCULE-5948770040. Values in parenthesis indicate highest resolution shell.**

### 5. MD simulations

**Fig. S4.**
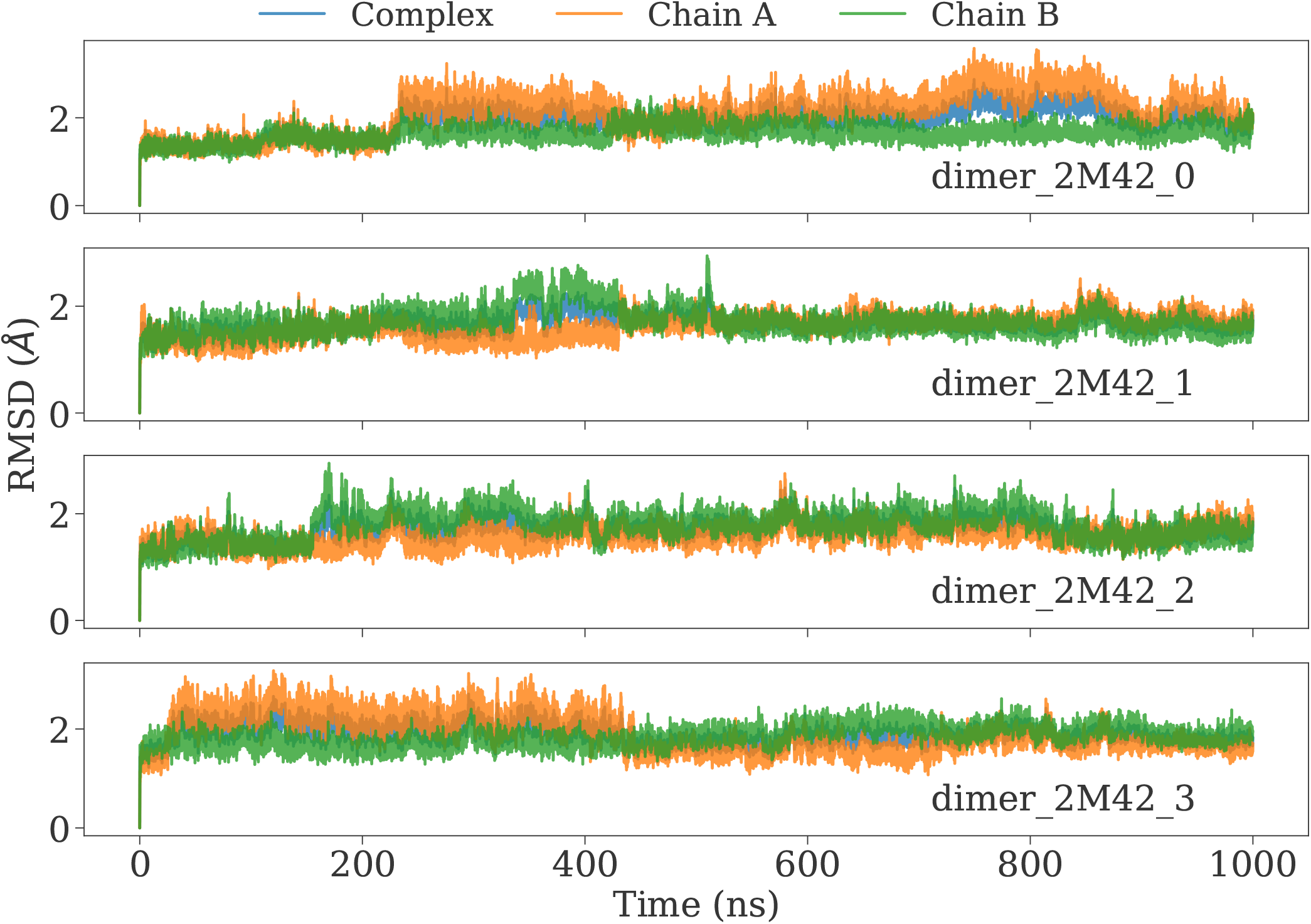
Root mean squared deviations from MD simulations of 2 MCULE-5948770040 molecules bound to each M^pro^ protomer. The RMSD for the complex is depicted in blue color whereas the individual protomers (chains A and B) are depicted in orange and green respectively.

#### ANCA-AE implementation details

In the methods section, we provided an overview of how ANCA-AE was implemented. Here we provide supporting information regarding its evaluation. In particular, ANCA-AE implements an autoencoder that takes as inputs the linear projections of the simulation datasets analyzed using ANCA. For our implementation, we noted that the optimal set of linear dimensions for ANCA with the LB and LF simulations (where the ligand was bound to both protomers) was 40, whereas including the simulations with the ligand bound to only one protomer, the optimal number of dimensions for ANCA was 50. We then mapped the autoencoder loss (based on the mean squared error in reconstructing the latent space) as a function of the number of latent dimensions (Fig. S7A), which showed that as we increased the number of latent dimensions, the loss value typically reduces. The other plots (Fig. S7B-D) summarize the characteristics of training and validation loss based on the model and the hyperparameters that were used in our training runs.

#### Analysis of MCULE-5948770040 bound to individual protomer units

We present an analysis of MCULE-5948770040 bound to one of the protomer units, rather than both protomers to check (1) if such a simulation would be stable, and (2) if the ligand bound to a particular protomer would induce the same long-range stability of the region R5 from our data. Our analysis here based on the RMSF profiles in Fig. S9A shows that compared to the ligand-free (LF) simulations, the RMSF of the ligand-bound (LB) states (either one or two ligands, indicated in the brackets) are significantly smaller. Thus, the presence of the ligand in either protomer can stabilize R5, apart from the loops in the binding region (identified as S2, S1, S4 and S3).

Further, the projections of these simulations using the ANCA-AE methods also illustrates that there are significant differences in the collective motions characterized by the LB and LF simulations (Fig. S9B and Fig. S11B). Notably, while there is some overlap between the LF and LB states in their collective motions, we observe that once the ligand binds to a particular chain, the fluctuations in LB-chain B are different fro LB-chain A as is often seen by the orientation of the ligand in the binding site. This illustrates that the conformational changes sampled by each of the LB-states are different and gives rise to unique set of conformational states sampled. These motions can be attributed to the additional flexibilty observed in the loops surrounding the ligand binding pocket(s) in the LF-state (and when the ligand is absent from one of the chains), giving rise to the collective motions that were shown in Fig. 4 of the main text.

**Fig. S5.**
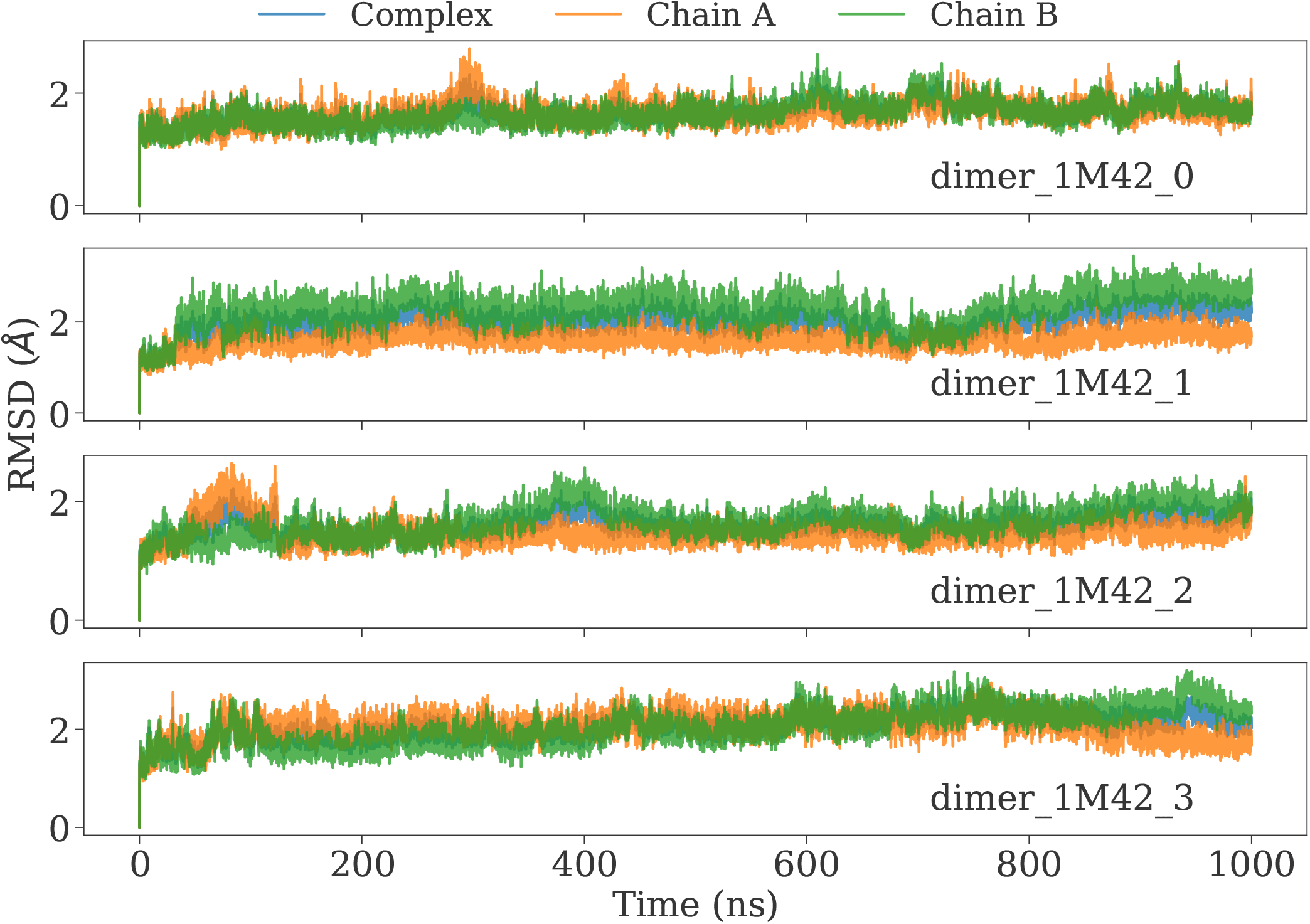
Root mean squared deviations from MD simulations of one MCULE-5948770040 bound to one of the M^pro^ protomer chains in the dimer. The RMSD for the complex is depicted in blue color whereas the individual protomers (chains A and B) are depicted in orange and green respectively.

### 6. Docking Pipeline Result Comparison

*Overlap between X-Chem fragments and MCULE-5948770040* We retrieved the fragments for 3CL-M^pro^ from X-Chem fragment crystalgraphic screens (4). Fragments did not have a high fingerprint similarity, with two common fragments appear in the highest 0.4 bin (see Fig. S12a). The crystals were aligned in PyMol and visualized in Fig. S12b.

**Fig. S6.**
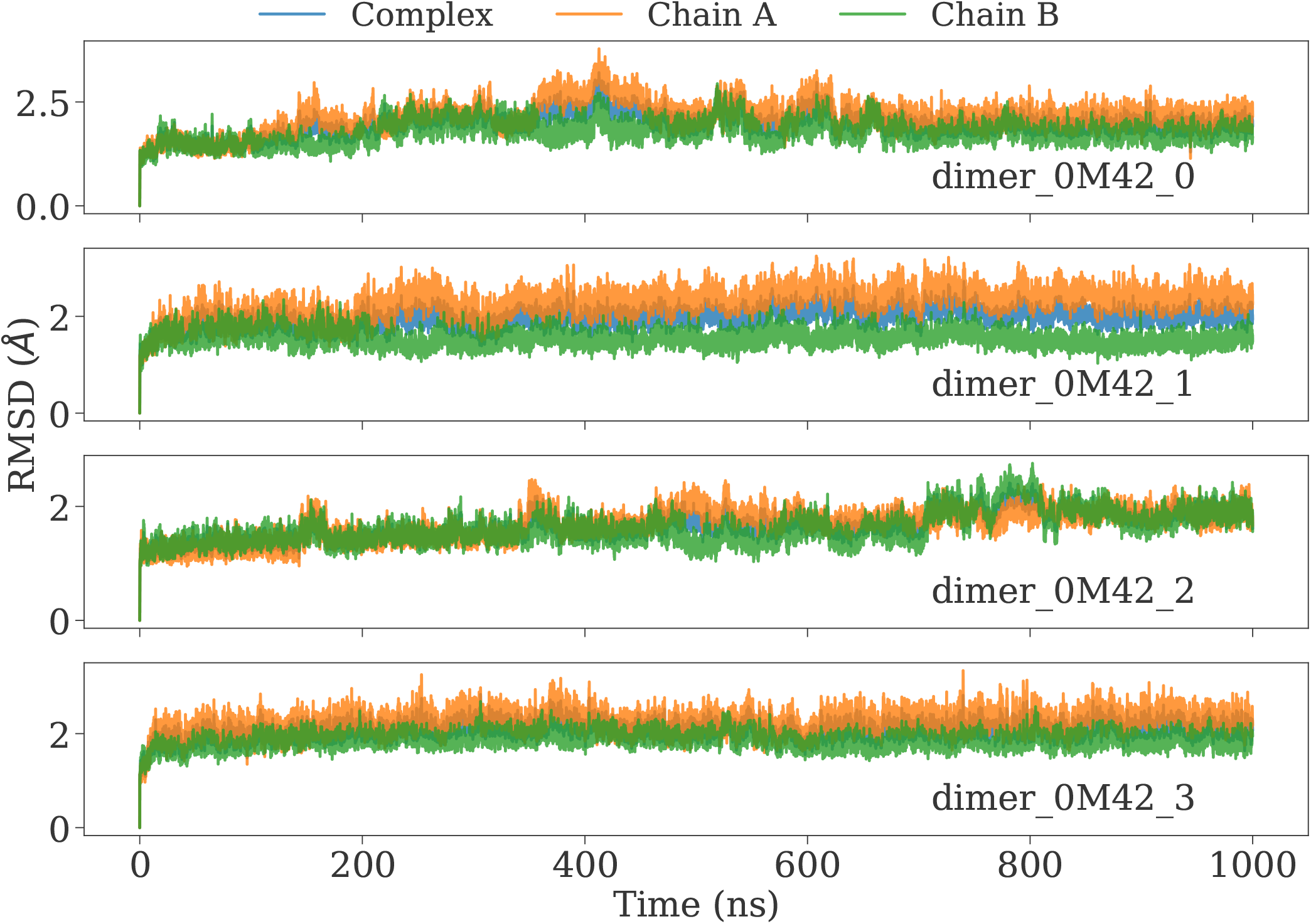
Root mean squared deviations from MD simulations of M^pro^ ligand-free state.

---

**Fig. S7.**
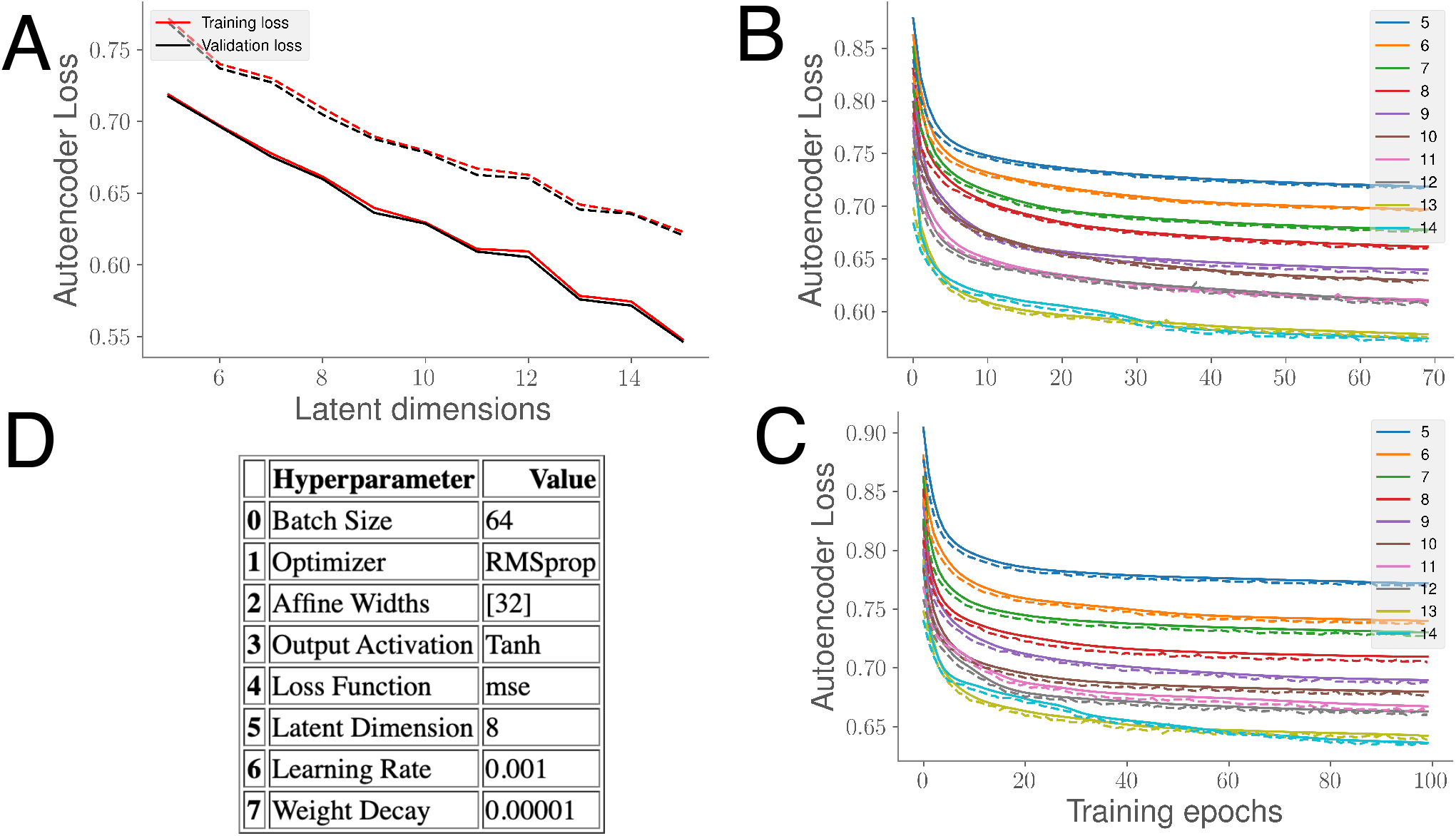
Characterizing how ANCA-AE identifies conformational substates from long timescale simulations. (A) Mapping the autoencoder loss versus the latent dimensions for training (red lines) and valiation (black) data based on 80:20 split of the simulation data. The solid lines represent the simulation datasets from the LB and LF states where the ligand was bound to both protomer units whereas the dotted line represents the loss based on including the simulations where the ligand was just bound to one of the protomers. (B) and (C) represent the training curves for ANCA-AE based on the number of latent dimensions (from 5-14) for both set of simulations as in (A). The solid lines track the training loss where as the dotted lines represent the validation loss as tracked by ANCA-AE. (D) summarizes the hyperparameter settings we used in training the ANCA-AE model.

**Fig. S8.**
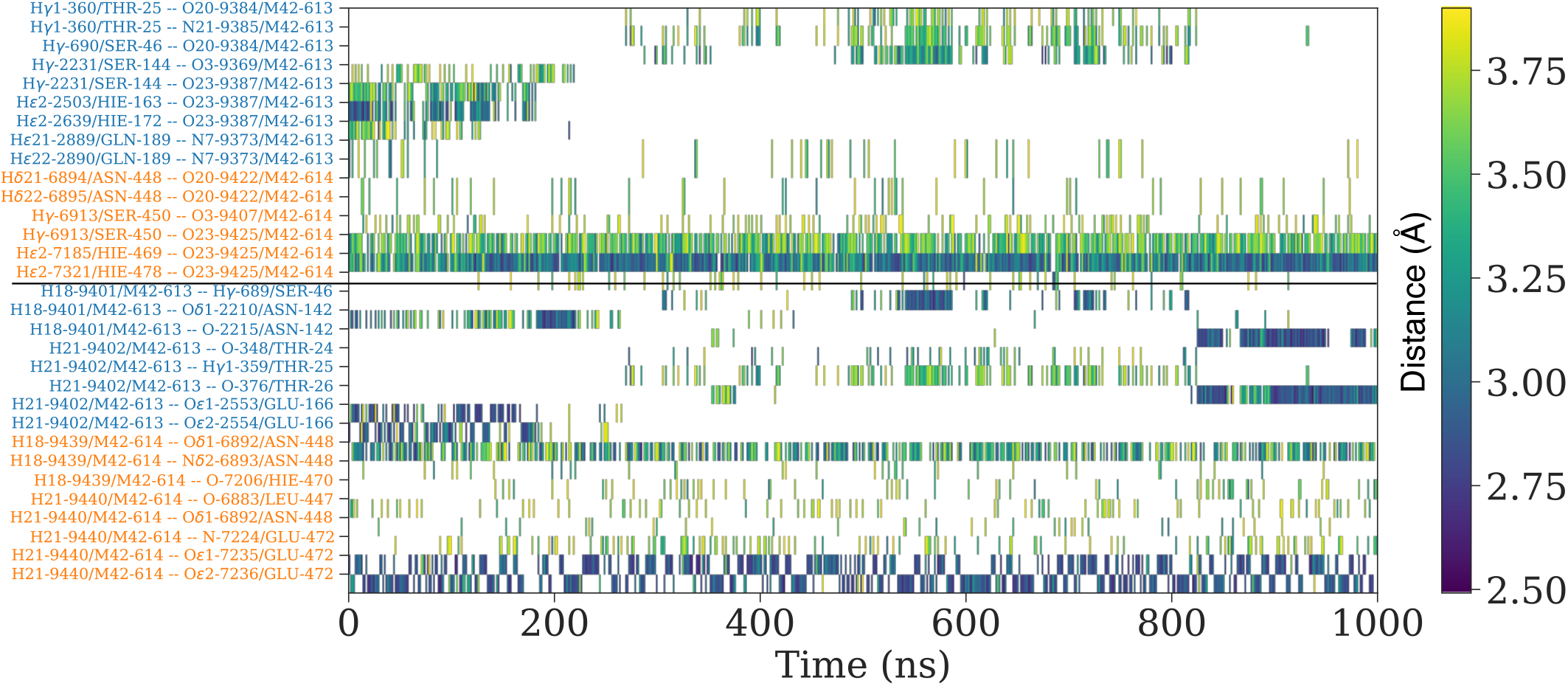
Hydrogen bond analysis in the LB simulations (bound to both protomers) show distinct behavior of the interaction patterns between the two protomers. The hydrogen bonding distance between the ligand and the protomer A is shown on the top with a solid line depicting the separation between the two protomers. While we observe stable hydrogen bonding patterns (distance between the heavy atoms < 2.75 Å) in protomer B, this is not observed in protomer A, where many hydrogen bonds are transient. Given that we observe a similar behavior with a second set of force-field parameters (calculated using NWChem), we posit that such fluctuations across the different protomers are unique.

**Fig. S9.**
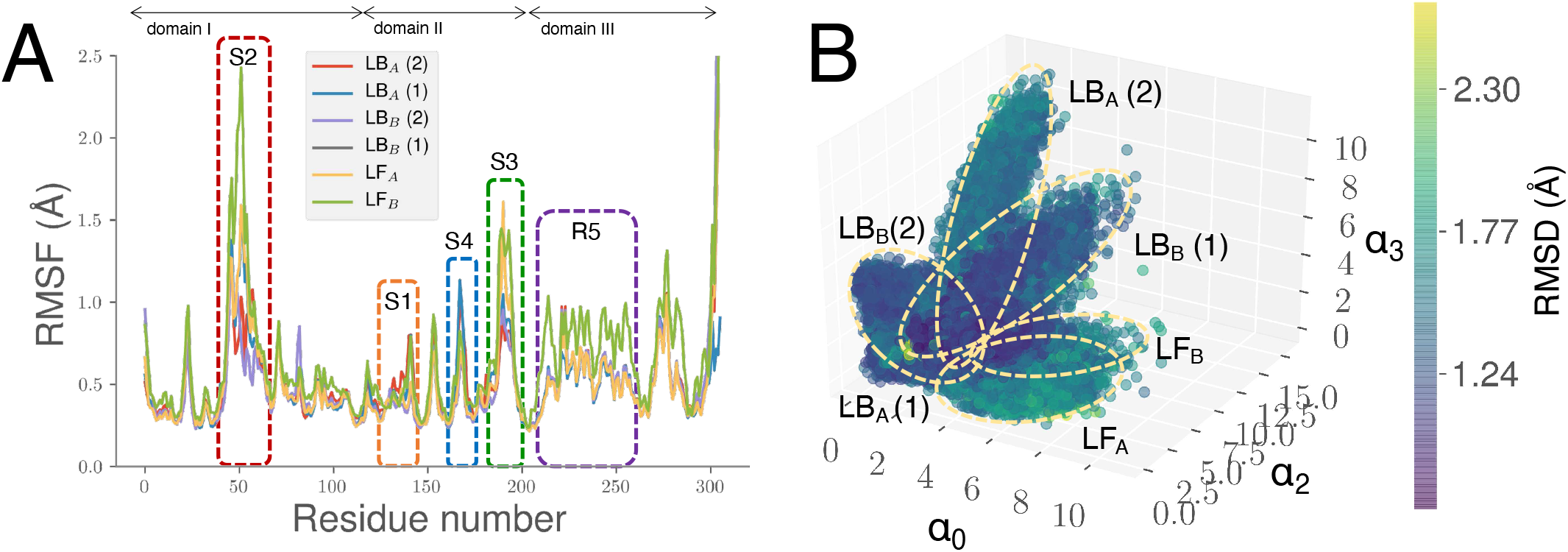
Conformational landscape of the LB-states as a consequence of binding MCULE-5948770040 to one of the protomers and its comparison to the other simulations. (A) Root mean squared fluctuations from MD simulations of M^pro^ the LB- and LF-states. The number in the brackets for the LB-states indicate the number of ligands bound to the complex. While we find similar regions affected by the ligand (Fig. 4 of the main paper), there are subtle differences in the overall profiles, indicating that a single molecule of MCULE-5948770040 bound to one of the protomers does not alter the overall conformational fluctuations as two molecules of the ligand bound to the complex. (B) Collective conformational fluctuations as characterized by ANCA-AE demonstrate the presence of distinct conformational states between the LB- and LF-states. The ellipses indicate a mixture of Gaussians taht depict how clusters are elucidated; further, simulations can be separated as shown in Fig. S11B.

**Fig. S10.**
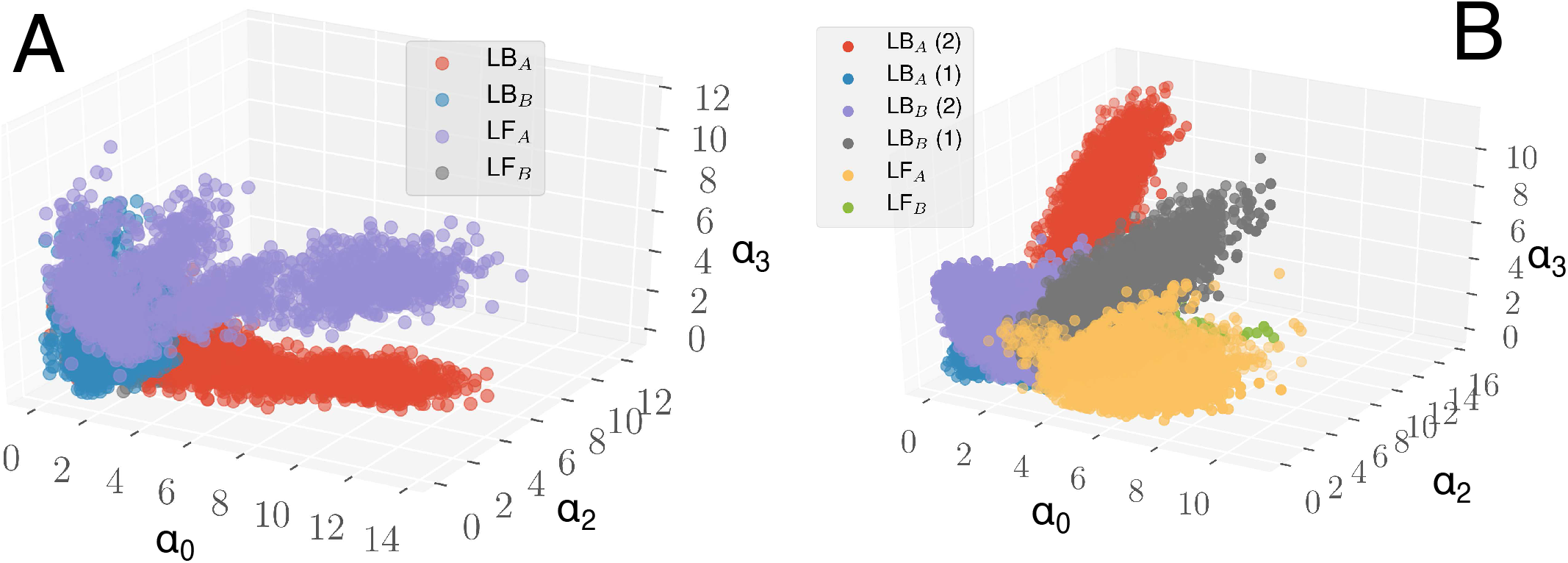
Conformational states sampled by the LB- and LF-states indicate distinct conformational transitions. (A) The projections from three ANCA-AE dimensions depicting distinct conformational transitions in the LB- and LF-states; notably the fluctuations in the LB-states in the two chains are quite distinct. (B) Same information as in (A), but includes projections from the two additional simulations where the ligand was bound only to one protomer units.

**Fig. S11.**
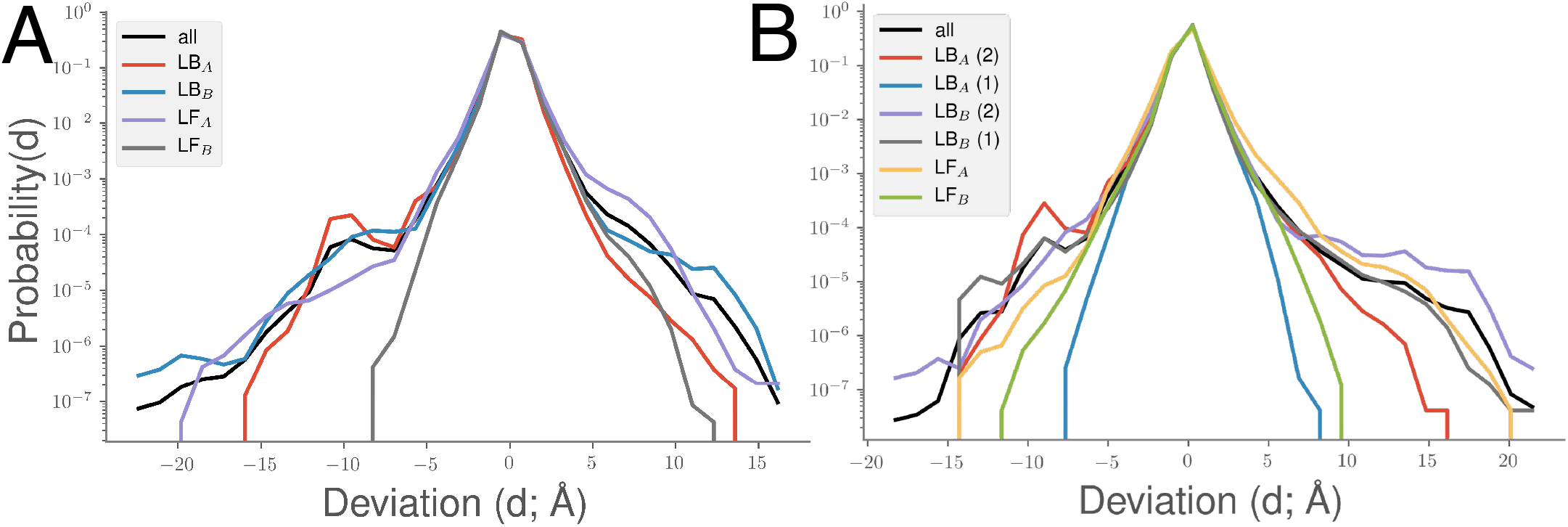
Presence of significant long-tails in the atomic fluctuations of the M^pro^ simulations, as quantified by the histograms of the positional deviations of the C^α^ atoms in our simulations. (A) depicts the long tails from the LB-(ligand bound to both protomers) and LF-states; (B) depicts the same information along with the simulations where the ligand was bound to only one of the two chains. Note that the fluctuations when including the single ligand bound to the protomer actually increases the overall fluctuations as observed from the deviation profiles.

**Fig. S12.**
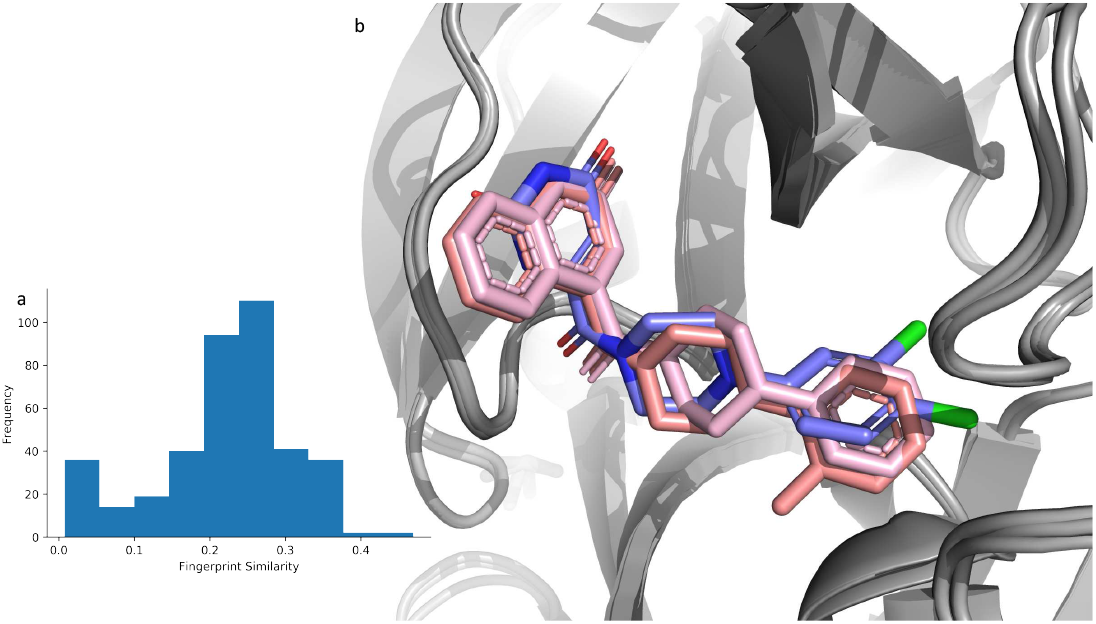
(a) Histogram outlining similarity of X-Chem 3CL-M^pro^ fragments with MCULE-5948770040 utilizing 2D fingerprint similarity. (b) X-Chem fragments x3303 and x11366 overlain with MCULE-5948770040 (4). There is an overlap on the P1 and linker, but notice the difference in the P2 region.

## Footnotes

* https://covid19.who.int

